# A learned oral signal drives long-lasting thirst satiation

**DOI:** 10.64898/2026.09.25.754368

**Authors:** Zhengya Liu, Ho Namkung, Jingkun Zhang, Julianne Davis, Naymalis La Santa Medina, Sophia Laskaris, Anyu Cao, Tianyi Chen, Xinlan Wen, Zachary A. Knight

## Abstract

Drinking quenches thirst long before the ingested water has been absorbed. Here we investigate the mechanism by combining sham drinking, which uncouples water ingestion from rehydration, with recordings of thirst-promoting neurons in the brain. We show that sham drinking suppresses thirst and rapidly inhibits thirst-promoting neurons despite no water absorption. Remarkably, this suppression of thirst neurons in the absence of rehydration is proportional to the amount of water consumed and persists for tens of minutes after drinking has stopped. This sustained inhibition is triggered by passage of water through the oral cavity, but not licking or swallowing alone, and undergoes extinction with repeated sham drinking, indicating it is learned. Complementary experiments using hypertonic saline demonstrate this learning is bidirectional. These findings reveal that the experience of rapid thirst quenching is not a hard-wired response to the consumption of water, but rather a learned prediction of future bodily state.

## Introduction

Internal needs are often experienced as localized bodily sensations. For example, thirst is caused by an increase in the osmolality of the blood^1–5^ but often experienced as a feeling of dryness in the mouth and throat^6^. How these localized somatic cues become coupled to general internal needs to guide motivated behaviors is not well understood.

For thirst, orosensory cues are important, in part, because there is a delay of many minutes between the ingestion of water and its absorption into the blood. Therefore, thirst-quenching must be driven by signals that precede water absorption and arise from either the mouth, throat, or GI tract^7,8^. These pre-systemic cues include a signal from the oropharynx and esophagus, which is thought to be short-lived^9^ and linked to either the taste of water^10^ or the act of swallowing^11^, and a gastrointestinal signal, which is thought to be slower and more durable^11–13^ (**Fig. 1A**). However, the fact that only a few seconds are required for the passage of water from the mouth to the intestine^14^ makes it challenging to disentangle these sequential cues and study them in isolation. As a result, despite decades of research^7^, it remains unclear what exactly each of these pre-systemic signals represents; how long each signal persists; and how they are functionally related to each other. One fundamental, but unanswered, question is whether these various thirst-quenching mechanisms are independent, hard-wired sensory signals, as is often assumed^15,16^, or whether they are instead organized in a hierarchy through learning.

**Figure 1.**
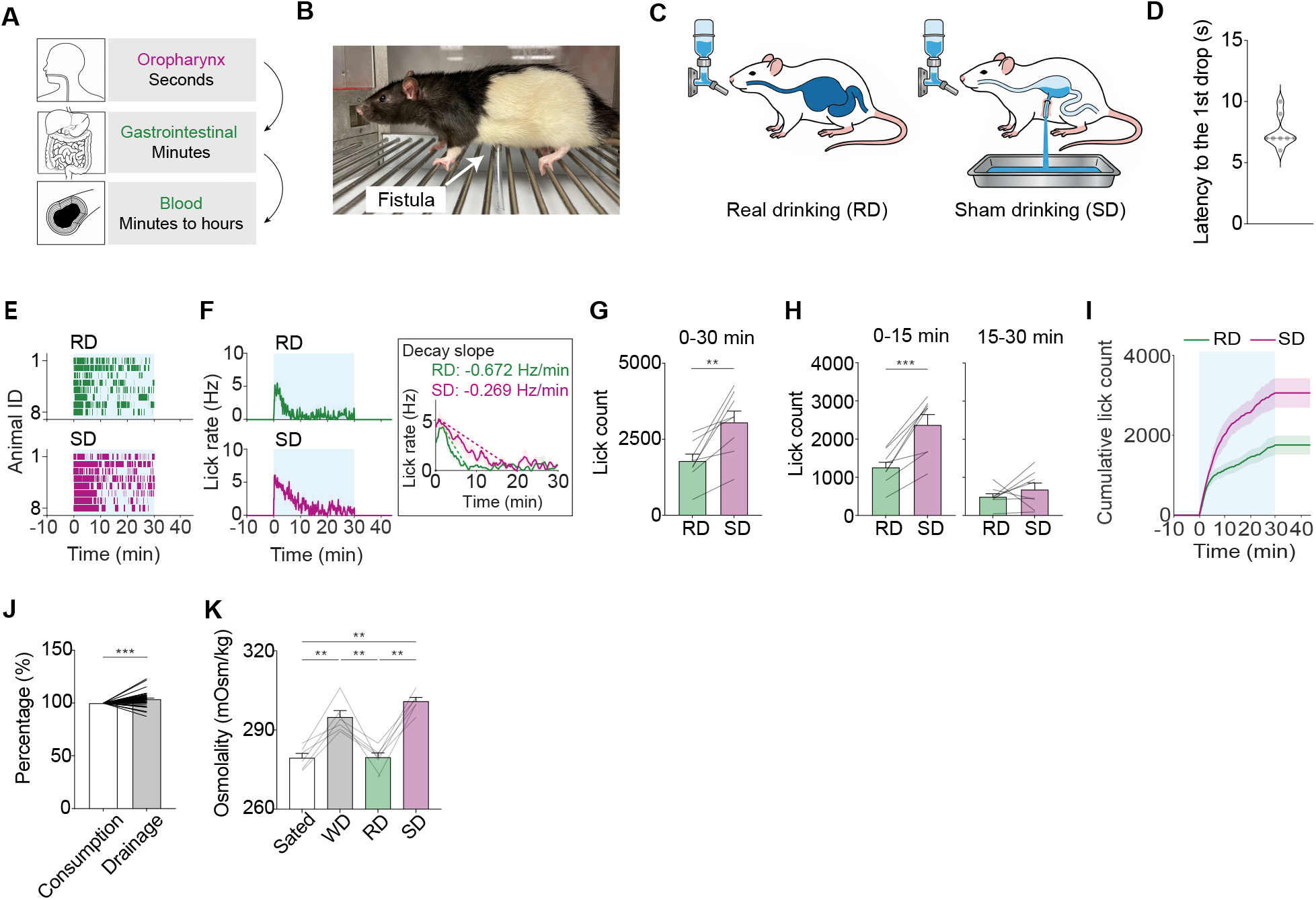
Sham drinking prevents water absorption but still leads to the cessation of drinking. **A,** Schematic illustrating candidate signals that contribute to thirst quenching across different timescales. **B,** Representative photograph of a rat implanted with a gastric fistula used for sham-drinking experiments. **C,** Schematic illustrating real drinking (RD), in which the gastric fistula is closed and ingested water is retained, and sham drinking (SD), in which the fistula is open and ingested water drains from the stomach. **D,** Latency from the first lick to the first drop of gastric drainage during SD. **E,** Lick rasters during the 30-min access period in RD and SD. **F,** Lick rate during the 30-min access period (left) and the corresponding decay slope (right) in RD and SD. **G,** Total lick count during the 30-min access period in RD and SD. **H,** Lick count during the first (0-15 min) and second (15-30 min) halves of the access period in RD and SD. **I,** Cumulative lick count during RD and SD. **J,** Water consumption and gastric drainage during SD, expressed as a percentage of total consumption. **K,** Blood osmolality measured 15 min after the session in sated, water-deprived (WD), RD, and SD conditions. Sample sizes: **D**, n = 8 sessions; **E-I**, n = 8 rats; **J**, n = 45 sessions; **K**, n = 6 rats. \**p* < 0.05, \*\**p* < 0.01, \*\*\**p* < 0.001. All values are reported as mean ± SEM. More detailed statistical information is in **Table S1**.

Answering these questions will require isolating individual pre-systemic signals and then measuring their contribution, in an awake, behaving animal, to thirst and its neural representation. We reasoned that this could be achieved by combining two techniques: sham drinking, which uncouples water ingestion from absorption and therefore isolates pre-gastric cues in freely behaving animals, and recording of thirst-promoting neurons in the brain, which reveals the contribution of these cues to thirst satiation quantitatively and in real-time.

## Results

### Sham drinking prevents water absorption but still leads to the cessation of drinking

To enable separation of pre- and post-gastric signals during natural ingestion, we equipped rats with a reversible gastric fistula (**Fig. 1B**). When the fistula is open (sham drinking, SD), ingested water drains out the stomach and thus no water absorption occurs (**Fig. 1C**). When the fistula is closed (real drinking, RD), water absorption is normal (**Fig. 1C**). We performed these studies in rats, rather than mice, because their larger size allows them to tolerate fistula implants for chronic experiments, which has not been demonstrated in smaller animals.

We first characterized drinking behavior in these animals. Rats were water-deprived for 24 hours, then given access to a sipper for self-paced water consumption. During real drinking, rats drank almost immediately when the sipper became available (**Fig. 1E, S1A;** latency, 5.7 ± 2.5 s), continued to drink at near maximum rate for 2 min, and then progressively decreased their drinking rate between the 2^nd^ and 8^th^ minute (**Fig. 1F**, −0.67 Hz/min). After the 8^th^ minute, further drinking was minimal (**Fig. 1F**), suggesting that the animals were satiated.

We next tested the same animals with the fistula open and measured their response to sham drinking for the first time. Similar to real drinking, rats drank voraciously as soon as the sipper became available (**Fig. 1E, S1A;** latency, 3.82 ± 1.21 s) and then showed a progressive decline in drinking rate (**Fig. 1F**). However, this rate of decline was slower during sham drinking (**Fig. 1F**; −0.269 Hz/min) and, as a result, animals consumed more water throughout the session (**Fig. 1G, I, Fig. S1B;** lick count: RD, 1757 ± 238; SD, 3059 ± 371; *p* = 0.0043; consumption: RD, 13.4 ± 1.3 mL; SD, 24.4 ± 2.7 mL; *p* = 0.0016). This increased intake was largely confined to the first 15 min (**Fig. 1H**; first 15 min: RD, 1263 ± 156 licks; SD, 2375 ± 272 licks; *p* = 0.00059), and, by the midpoint of the 30 min session, rats had reduced their rate of sham drinking to the level of real drinking controls (**Fig. 1H**; second 15 minutes: RD, 494 ± 90 licks; SD, 684 ± 169 licks; *p* = 0.37).

The fact that rats progressively reduce their rate of sham drinking suggests they experience satiation without water absorption, although other explanations are possible. One alternative explanation is that the rats became physically exhausted. However, this was not the case because we showed that hungry rats would drink nearly twice as much fluid during a 30 min sham feeding test with sucrose (**Fig. S1C-E;** lick count: SD, 3059 ± 371; SF, 5668 ± 397; *p* = 0.00028; consumption: SD, 24.4 ± 2.7 mL; SF, 41.8 ± 3.4 mL; *p* = 0.0013). A second possibility is that some water bypassed the fistula and was absorbed, resulting in rehydration. However, we confirmed that this did not occur through three analyses. First, we visually observed that the ingested fluid drained from the fistula with short latency (**Video S1, Fig. 1D**, time from the first lick to the first drop of drainage 7.5 ± 0.5 s), which we confirmed by simultaneous X-ray fluoroscopy (**Video S2**). Second, we collected the fluid that drained from the fistula and compared this to the volume of fluid that was consumed. This revealed that drainage exceeded intake (**Fig. 1J**; 104% of intake; *p* = 0.00045), likely due to the contribution of small amounts of saliva and gastric fluids. This indicates that little, if any, ingested water failed to drain through the fistula. Third, we measured the blood osmolality 15 min after each session concluded (45 min after the start of drinking). As expected, water-deprived (WD) rats had a significantly elevated blood osmolality at the start of the session (**Fig. 1K**; Sated, 279.4 ± 1.7 mOsm/kg; WD, 294.8 ± 2.5 mOsm/kg; multiple-comparison test, Sated vs. WD, *p* = 0.001) and this returned to baseline after real drinking (RD, 279.6 ± 1.8 mOsm/kg; Sated vs. RD, *p* = 0.90). In contrast, there was no decrease in blood osmolality after sham drinking sessions (**Fig. 1K**; SD, 300.82 ± 1.54 mOsm/kg; WD vs. SD, *p* = 0.13; RD vs. SD, *p* = 0.001), indicating systemic rehydration did not occur. Together, these data suggest that sham drinking rats experience delayed, but otherwise normal, thirst satiation despite the lack of any water absorption or rehydration.

### MnPO^Agtr1a^ neurons drive drinking and recapitulate the hallmarks of thirst

To probe the mechanisms that drive satiation during sham drinking, we investigated the role of the median preoptic nucleus (MnPO), which is the key integration site for thirst in the brain^17^. In the mouse, thirst-promoting MnPO neurons are labeled by expression of the angiotensin receptor 1a (*Agtr1a*)^18^. To gain genetic access to these cells in the rat, we generated a knock-in line that expresses Cre recombinase from the *Agtr1a* gene (Agtr1a-T2A-Cre: **Fig. 2A**) and confirmed this line induces a recombination pattern that mirrors endogenous *Agtr1a* expression (**Fig. 2B**).

**Figure 2.**
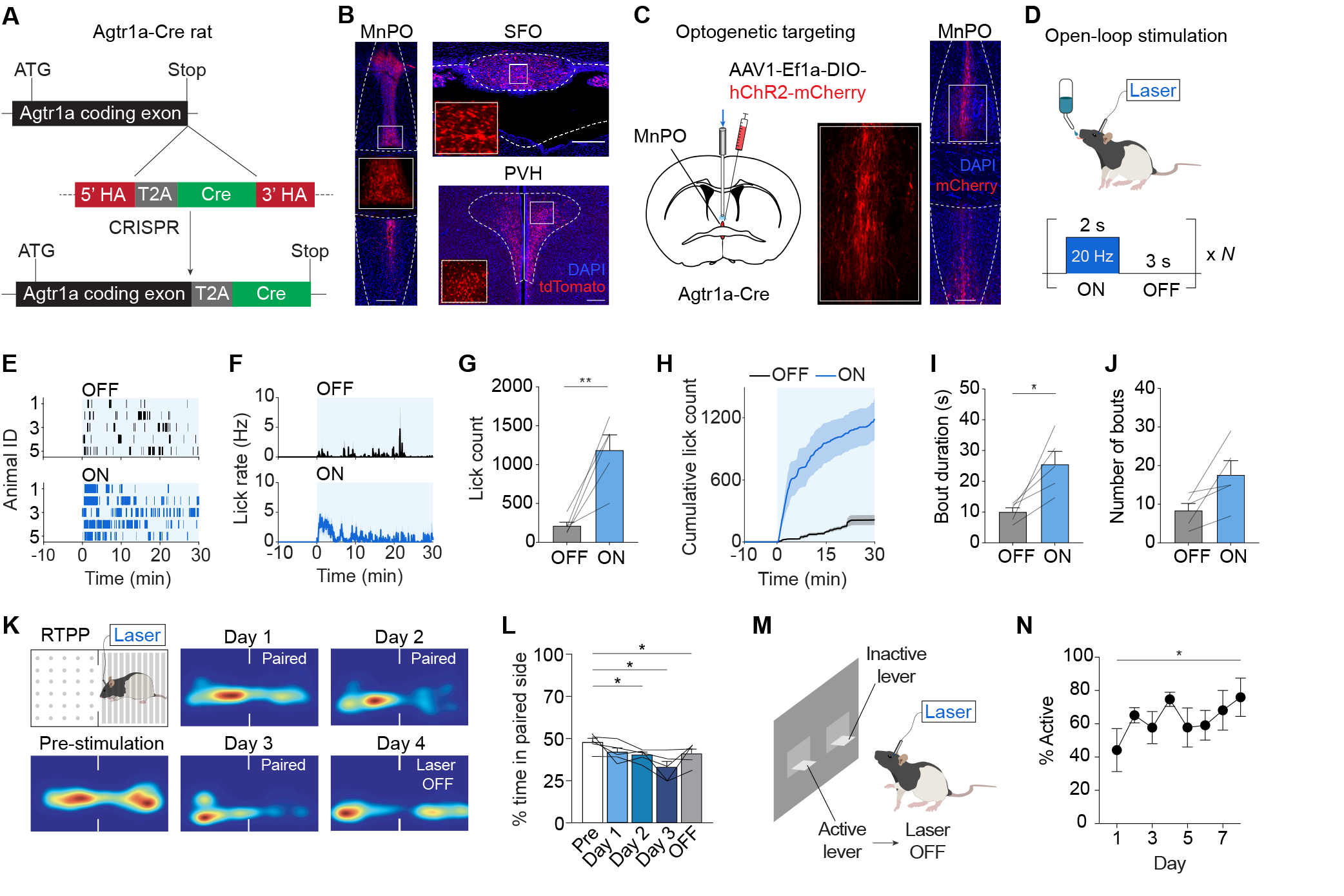
Genetic access and functional validation of thirst neurons in the MnPO. **A,** Generation of the Agtr1a-T2A-Cre knock-in rat by CRISPR-mediated homologous recombination. **B,** Agtr1a reporter expression in the median preoptic nucleus (MnPO), subfornical organ (SFO), and paraventricular hypothalamus (PVH) in Agtr1a-T2A-Cre rats crossed with a tdTomato reporter line. **C,** Optogenetic targeting of the MnPO: schematic of the AAV injection site and optic-fiber placement (left) and representative ChR2-mCherry expression in the MnPO (right). **D,** Open-loop optogenetic stimulation protocol during access to a water bottle. **E,** Lick rasters from sated rats during 30-min water access in laser-OFF and laser-ON sessions. **F,** Lick rate during laser-OFF and laser-ON sessions. **G,** Total lick count in laser-OFF and laser-ON sessions. **H,** Cumulative lick count in laser-OFF and laser-ON sessions. **I,** Average drinking-bout duration in laser-OFF and laser-ON sessions. **J,** Number of drinking bouts in laser-OFF and laser-ON sessions. **K,** Real-time place-preference (RTPP) assay for optogenetic stimulation, with representative occupancy heatmaps during pre-stimulation, laser-paired sessions (Days 1-3), and a subsequent laser-OFF session (Day 4). **L,** Percentage of time spent on the laser-paired side across RTPP sessions. **M,** Schematic of the lever-pressing task used to test reinforcement by optogenetic stimulation. **N,** Percentage of active-lever presses among total lever presses across training days. All scale bars are 200 µm. Sample size: **E-N,** n = 5 rats. \**p* < 0.05, \*\**p* < 0.01. All values are reported as mean ± SEM. More detailed statistical information is in **Table S1**.

We first tested whether stimulation of MnPO^Agtr1a^ neurons would drive drinking. We injected a Cre-dependent AAV expressing channelrhodopsin-2 into the MnPO of *Agtr1a^Cre^* rats and then installed an optical fiber above the injection site (**Fig. 2C**). We found that laser stimulation (20 Hz) caused voracious water consumption in sated rats (**Fig. 2D-H, Fig. S2B;** lick count: OFF, 217 ± 49; ON, 1187 ± 196; *p* = 0.0075; consumption: OFF, 1.5 ± 0.2 mL; ON, 10.1 ± 2.3 mL; *p* = 0.017). This occurred through an increase in bout duration (**Fig. 2I**; OFF, 10.1 ± 1.4 s; ON, 25.6 ± 4.1 s; *p* = 0.022) and a trend toward an increase in bout number (**Fig. 2J**; OFF, 8.4 ± 1.9; ON, 17.6 ± 3.7; *p* = 0.055). There was also a trend toward a shorter latency to the first lick (**Fig. S2A;** OFF, 49.8 ± 13.7 s; ON, 19.6 ± 6.4 s; *p* = 0.13). On the other hand, there was no effect of MnPO^Agtr1a^ stimulation on food intake (**Fig. S2J;** OFF, 2.5 ± 0.6 g; ON, 2.1 ± 0.6 g; *p* = 0.64). Thus, stimulation of MnPO^Agtr1a^ neurons specifically drives drinking.

MnPO neurons are thought to encode the negative sensation of thirst^18,19^. To test this, we first measured the valence of MnPO^Agtr1a^ neuron activation using a real-time place preference (RTPP) assay. Rats were allowed to move between two chambers with distinct visual cues (wall patterns, dot vs stripe) and received laser stimulation only on the paired side (**Fig. 2K**). During a baseline test with no laser, rats showed no preference for either side (**Fig. 2K-L**; 47.8 ± 2.3% time on the paired side; *p* = 0.40 versus 50%). However, during subsequent days of testing with laser stimulation, animals developed a clear preference for the unpaired side, indicating that laser stimulation has negative valence (**Fig. 2K-L**; one-way repeated-measures ANOVA, *p =* 0.002; multiple-comparison test versus baseline: Day 1, *p* = 0.35; Day 2, *p* = 0.025; Day 3, *p* = 0.022). This valence persisted when the rats were tested in extinction (Laser OFF), indicating that they had learned to associate one side of the chamber with an aversive state (**Fig. 2K-L**; OFF, 41.0 ± 2.9%; multiple-comparison test versus baseline, *p* = 0.014).

Actions that promote relief from an aversive state are learned through negative reinforcement, and this is an important mechanism that promotes drinking^18,19^. To test whether MnPO^Agtr1a^ neuron activity is negatively reinforcing, we allowed rats to press a lever to pause (1 min) optogenetic stimulation of MnPO^Agtr1a^ neurons (**Fig. 2M**). During a 4-hour session across multiple days, rats learned to preferentially press the active lever to pause laser stimulation (**Fig. 2N**; one-way repeated-measures ANOVA, *p* = 0.042; multiple-comparison test, Day 1 vs. Day 8, *p* = 0.012). Together, these data show that stimulation of MnPO^Agtr1a^ neurons drives drinking, has negative valence, and supports learning through negative reinforcement, indicating that these cells can recapitulate key hallmarks of thirst^18–20^.

### Sham drinking causes rapid and sustained inhibition of MnPO^Agtr1a^ neurons

We next investigated the activity dynamics of MnPO^Agtr1a^ neurons during real and sham drinking. We injected a Cre-dependent AAV expressing GCaMP6s into the MnPO of *Agtr1a^Cre^*rats and installed an optical fiber above the injection site for calcium recordings by fiber photometry (**Fig. 3A**). In a second surgery, rats were equipped with a reversible gastric fistula for sham drinking. To confirm that MnPO^Agtr1a^ neurons can sense changes in bodily hydration, we administered an intraperitoneal (IP) injection of hypertonic saline (5 M, 2 mL), which rapidly increases the osmolality of the blood^21^. This caused dramatic and sustained MnPO^Agtr1a^ neuron activation (**Fig. 3B**; 9.4 ± 2.1 z vs. −0.71 ± 0.38 z; 5 M NaCl vs. PBS control injection; *p* = 0.002), showing that these cells robustly respond to increases in blood osmolality.

**Figure 3.**
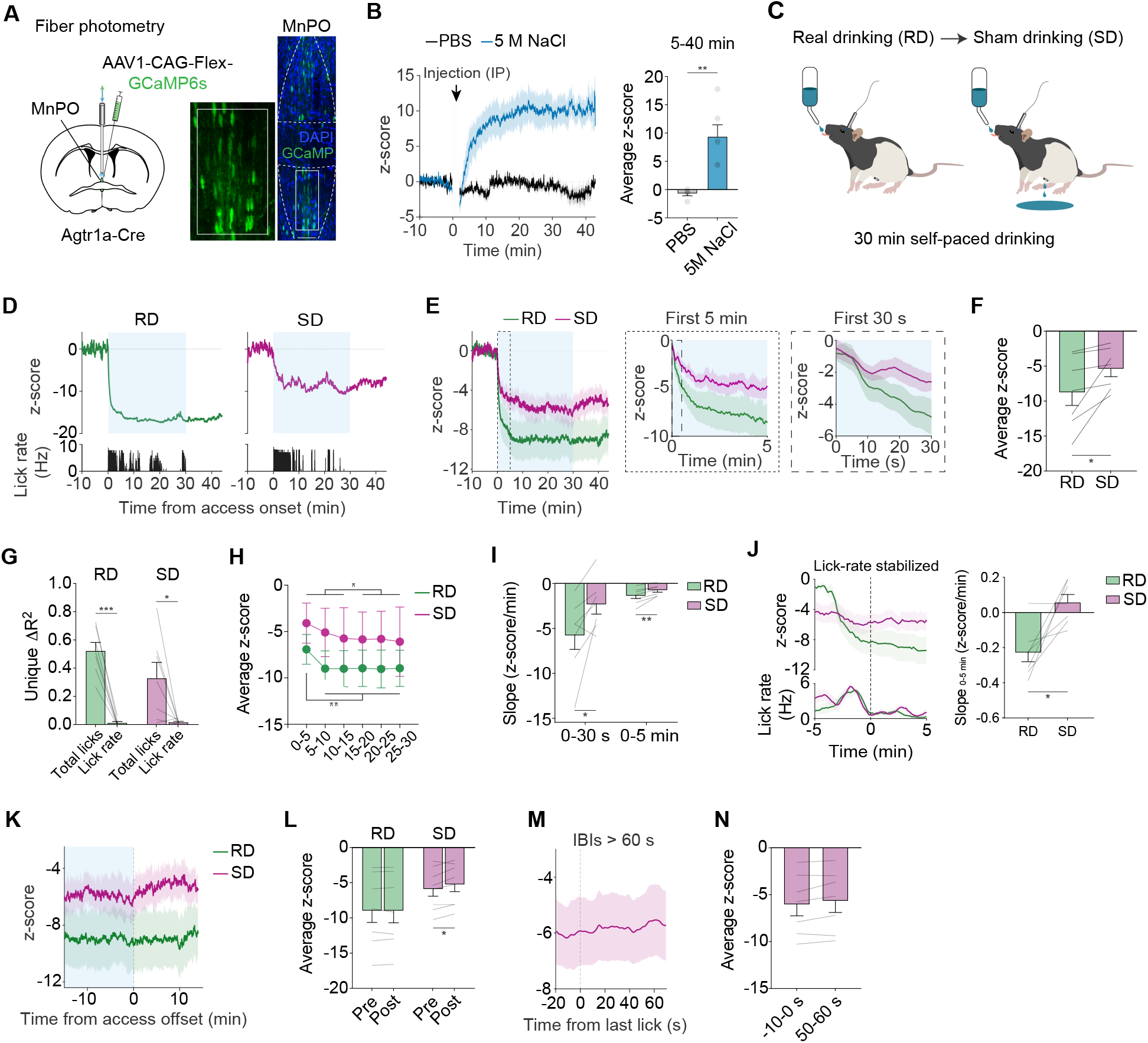
Pre-gastric signals strongly contribute to the rapid quenching of thirst. **A,** Fiber photometry recording of MnPO^Agtr1a^ neurons: schematic of the AAV injection site and optic-fiber placement (left) and representative GCaMP6s expression in the MnPO (right). **B,** Average MnPO^Agtr1a^ activity traces following intraperitoneal (IP) injection of PBS or 5 M NaCl (left) and mean z-score from 5-40 min after injection (right). **C,** Experimental timeline for 30-min self-paced water access during real drinking (RD) and sham drinking (SD) with fiber photometry recording. **D,** MnPO^Agtr1a^ activity (top) and lick rate (bottom) from a representative rat during RD and SD, aligned to access onset. Blue shading indicates the 30-min water-access period. **E,** Average MnPO^Agtr1a^ activity traces during RD and SD across the full session (left), with expanded views of the first 5 min (middle) and first 30 s (right) after access onset. **F,** Average MnPO^Agtr1a^ activity during RD and SD sessions. **G,** Unique variance in MnPO^Agtr1a^ activity (ΔR²) explained by cumulative lick count or instantaneous lick rate in a multivariable regression model, shown separately for RD and SD. **H,** Average MnPO^Agtr1a^ activity during consecutive 5-min epochs of the 30-min access period in RD and SD. For RD, the first 5-min epoch was compared with the average of the remaining five epochs; for SD, the average of the first three epochs was compared with the average of the last three epochs. **I,** Slope of the decline in MnPO^Agtr1a^ activity during the first 30 s and first 5 min after access onset in RD and SD. **J,** Average traces of MnPO^Agtr1a^ activity (top) and lick rate (bottom) aligned to the time at which lick rate stabilized at a low level (left), and the slope of MnPO^Agtr1a^ activity over the subsequent 0-5 min in RD and SD (right). **K,** Average MnPO^Agtr1a^ activity traces during RD and SD aligned to the end of the 30-min access period. **L,** Average MnPO^Agtr1a^ activity over approximately 15-min periods before and after access offset for the traces shown in **(K)**. **M,** Average MnPO^Agtr1a^ activity trace during SD aligned to the last lick of bouts followed by an inter-bout interval (IBI) >60 s. **N,** Average MnPO^Agtr1a^ activity from −10 to 0 s and from 50 to 60 s relative to the last lick for the bouts analyzed in **(M)**. Sample sizes: **B,** PBS n = 5 rats and 5 M NaCl n = 6 rats; **D-N,** n = 7 rats. \**p* < 0.05, \*\**p* < 0.01, \*\*\**p* < 0.001. All values are reported as mean ± SEM. More detailed statistical information is in **Table S1**.

To characterize how MnPO^Agtr1a^ neurons respond to water consumption, we water-deprived rats for 24 h and then gave them access to a sipper for self-paced, real drinking (30 min) (**Fig. 3C**). As expected^12,19^, MnPO^Agtr1a^ neurons were rapidly and strongly inhibited by drinking (**Fig. 3D-F)**, with activity dropping sharply upon water access and continuing to decline over the next several minutes as animals drank (**Fig. 3D-E**). This decrease in the activity of MnPO^Agtr1a^ was much better explained by cumulative water consumption than instantaneous lick rate (**Fig. 3G**; unique ΔR², 0.52 ± 0.06 vs. 0.013 ± 0.009; *p* = 0.00012). Within 5-10 minutes of water access, most animals had stopped drinking, and, accordingly, MnPO^Agtr1a^ neuron activity stabilized at a much lower level (**Fig. 3H**; 0–5 min vs. average 5–30 min, *p* = 0.0028; no further change from 5–30 min, one-way repeated-measures ANOVA, *p* > 0.99). These observations support a model^12,19^ in which the progressive inhibition of MnPO^Agtr1a^ neurons during drinking represents the satiation of thirst.

We next investigated how these neurons respond to drinking when the fistula is open and therefore no water is absorbed. Rats were water-deprived for 24 h and then given access to a sipper for self-paced, sham drinking (30 min) (**Fig. 3C**). Remarkably, sham drinking caused a rapid inhibition of MnPO^Agtr1a^ neurons that resembled real drinking in many respects. In both cases, MnPO^Agtr1a^ neurons were progressively inhibited over several minutes as rats drank (**Fig. 3E**). Like real drinking, the magnitude of inhibition during sham drinking was better explained by cumulative intake than by instantaneous lick rate (**Fig. 3G**; unique ΔR², 0.33 ± 0.11 vs. 0.016 ± 0.007; *p* = 0.037). The rate of this inhibition was slower during sham drinking (**Fig. 3I**; 0–5 min slope, RD, −1.4 ± 0.3 z/min; SD, −0.8 ± 0.2 z/min; *p* = 0.0044) and, accordingly, rats drank for longer before stopping and achieved less total inhibition (**Fig. 3F**; RD, −8.75 ± 1.92 z; SD, −5.38 ± 1.12 z; *p* = 0.036). Finally, during sham drinking, MnPO^Agtr1a^ activity reached a sustained inhibitory plateau immediately before lick rate declined and stabilized at a low level (**Fig. 3J**), suggesting that this decline in MnPO^Agtr1a^ neuron activity is causally involved in the cessation in drinking. During real drinking, on the other hand, MnPO^Agtr1a^ activity continued to decrease even after lick rate had stabilized at a low level, indicating that subsequent GI or post-absorptive signals can further inhibit MnPO^Agtr1a^ neurons. Taken together, these data indicate that pre-gastric signals alone are sufficient to reduce MnPO^Agtr1a^ neuron activity below the level necessary for the cessation of drinking.

Two aspects of the response of MnPO^Agtr1a^ neurons to sham drinking were particularly unexpected. The first is that the inhibition was progressive, meaning that it increased over minutes as animals continued to drink (**Fig. 3E**) and was strongly associated with cumulative water intake (**Fig. 3G**). This cannot be due to sensing of water in the GI tract^11–13^, which is absent in sham drinking, and indicates instead that MnPO^Agtr1a^ neurons can integrate over time the volume of water that passes through the mouth. The second surprising result is that the inhibition was stable. For example, we found that MnPO^Agtr1a^ neurons remained durably inhibited even when rats paused sham drinking for >1 min, (**Fig. 3M-N**; −10 to 0 s, −6.1 ± 1.2 z; 50 to 60 s, −5.7 ± 1.2 z; −10 to 0 vs. 50 to 60 s, *p* = 0.064; 50 to 60 s vs. baseline, *p* = 0.0038). Indeed, when water access was removed at the end of the 30 min trial, MnPO^Agtr1a^ neuron activity remained strongly inhibited relative to baseline for the remaining 15 min of the session (**Fig. 3K-L**; SD post-offset, −5.3 ± 1.2 z, *p* = 0.0047 vs. baseline). This indicates that pre-gastric signals alone can produce sustained inhibition of MnPO^Agtr1a^ neurons that long outlasts the transient sensation of water passing through the oral cavity.

### Pre-gastric water sensation is critical for the strong and sustained inhibition of thirst

We next sought to clarify the nature of the pre-gastric signal that drives the strong and sustained inhibition of MnPO^Agtr1a^ neurons. We reasoned that sham drinking involves at least three distinct components: licking, swallowing, and chemosensory detection of water in the mouth and/or esophagus (**Fig. 4A**). Consistent with previous work^20^, air licking at an empty sipper produced a rapid initial inhibition of MnPO^Agtr1a^ activity while animals licked vigorously (**Fig. 4B**). However, as licking rapidly declined over the subsequent period, MnPO^Agtr1a^ activity returned toward baseline (**Fig. 4B**). This contrasted with water sham drinking, during which activity remained suppressed over the same period (**Fig. 4E–F, I**; 30 s–5 min Δz-score: water SD, −2.1 ± 0.7 z; air licking, 1.3 ± 0.3 z; multiple-comparison test, water SD vs. air licking, *p* = 0.0009; 30 s–5 min Δz-score per 1,000 licks: water SD, −1.5 ± 0.5; air licking, 4.0 ± 1.1; multiple-comparison test, water SD vs. air licking, *p* = 0.0035). Thus, licking alone can produce a weak initial inhibition but is insufficient to generate the sustained response observed during water ingestion.

**Figure 4.**
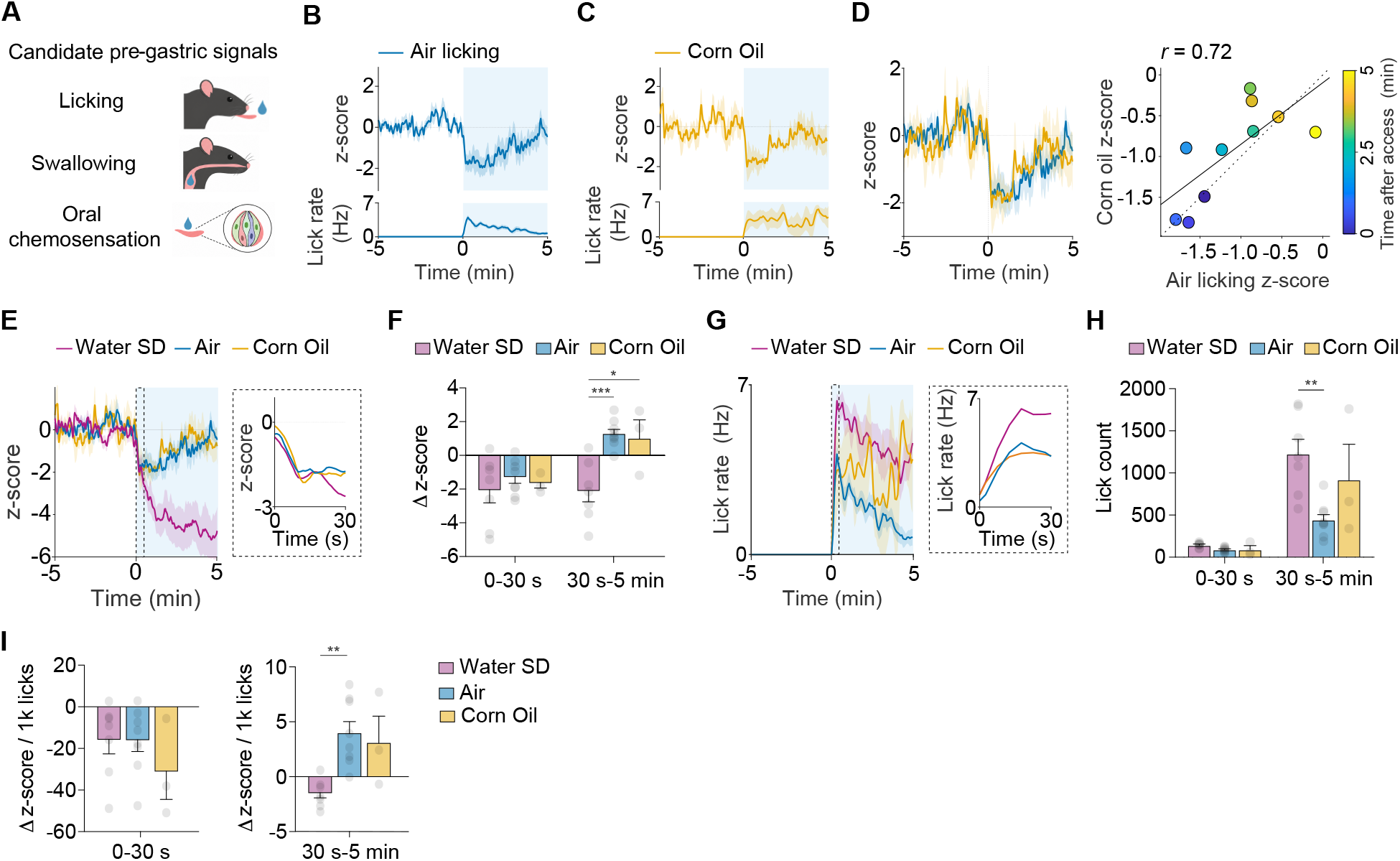
Pre-gastric water sensation is critical for the strong and sustained inhibition of thirst. **A,** Candidate components of pre-gastric signaling during thirst quenching. **B,** Average traces of MnPO^Agtr1a^ activity (top) and lick rate (bottom) during air licking from an empty bottle. **C,** Average traces of MnPO^Agtr1a^ activity (top) and lick rate (bottom) during corn-oil sham drinking. **D,** Overlay of average MnPO^Agtr1a^ activity traces during air licking and corn-oil sham drinking (left), and the relationship between their mean neural responses across ten consecutive 30-s bins during the first 5 min after access onset (right). Points are colored by time after access onset. **E,** Average MnPO^Agtr1a^ activity traces during the first 5 min after access onset across water sham drinking (SD), air licking, and corn-oil sham drinking (left), with an expanded view of the first 30 s (right). **F,** Change in MnPO^Agtr1a^ activity (Δz-score) during 0-30 s and 30 s-5 min after access onset, calculated as z(30 s) − z(0 s) and z(5 min) − z(30 s), respectively. **G,** Average lick-rate traces during the first 5 min after access onset across conditions (left), with an expanded view of the first 30 s (right). **H,** Lick count during 0-30 s and 30 s-5 min after access onset. **I,** Change in MnPO^Agtr1a^ activity (Δz-score) normalized by lick count within the same interval, shown separately for 0-30 s and 30 s-5 min and expressed as Δz-score per 1,000 (1k) licks. Sample sizes: water SD, n = 7 rats; air licking, n = 8 rats; corn-oil sham drinking, n = 3 rats. \**p* < 0.05, \*\**p* < 0.01, \*\*\**p* < 0.001. All values are reported as mean ± SEM. More detailed statistical information is in **Table S1**.

We next asked whether swallowing could account for this sustained inhibition. Rats readily sham drank corn oil, producing vigorous licking and swallowing (**Fig. 4G-H**) and, like air licking, corn-oil ingestion rapidly inhibited MnPO^Agtr1a^ activity during the initial phase (**Fig. 4C**). However, although animals continued to lick corn oil vigorously during the subsequent period, MnPO^Agtr1a^ activity nevertheless returned toward baseline (**Fig. 4C, E–H**). Accordingly, the temporal profile of the neural response to corn oil closely resembled that evoked by air licking (**Fig. 4D**; 10 time bins, Pearson’s *r* = 0.72, *p* = 0.020), and both differed from the sustained inhibition observed during water sham drinking (**Fig. 4E–F, I**; 30 s–5 min Δz-score: corn oil, 1.1 ± 1.1 z; multiple-comparison test, SD vs. corn oil, *p* = 0.0165; Δz-score per 1,000 licks: water SD, −1.5 ± 0.5; corn oil, 3.2 ± 2.4; multiple-comparison test, SD vs. corn oil, *p* = 0.0609). These findings indicate that neither swallowing nor sustained licking is sufficient to maintain strong MnPO^Agtr1a^ inhibition. Together, these results reveal two components of the pre-gastric response: a rapid initial inhibition that can be evoked by licking and swallowing, and a sustained inhibition that is selectively maintained during water ingestion. Thus, the strong and prolonged inhibition of MnPO^Agtr1a^ neurons requires the chemosensory detection of water in the mouth and/or esophagus.

### Inhibition of MnPO^Agtr1a^ neurons attenuates with repeated sham drinking experiences

The inhibition of MnPO^Agtr1a^ neurons during sham drinking is progressive (**Fig. 3E**) and durable (**Fig. 3K-N**) whereas the sensation of water passing through the mouth is transient. This indicates that the inhibition of MnPO^Agtr1a^ neurons during drinking is not merely a sensory response to an oral cue. Rather, it resembles a prediction of what that cue means for the subsequent restoration of fluid balance. If this is true, then the inhibition of MnPO^Agtr1a^ neurons during sham drinking should become attenuated as animals learn that drinking no longer predicts bodily rehydration.

To test this, we exposed rats to sham drinking across multiple days and measured how the neural response to water ingestion changed with experience. For these experiments, we used a trial-based paradigm in which rats had access to water in 5 min blocks separated by 2 min intervals in which the sipper was retracted (**Fig. 5A)**. This design allowed us to characterize the ingestion-linked and post-access neural responses separately at multiple timepoints within each session.

**Figure 5.**
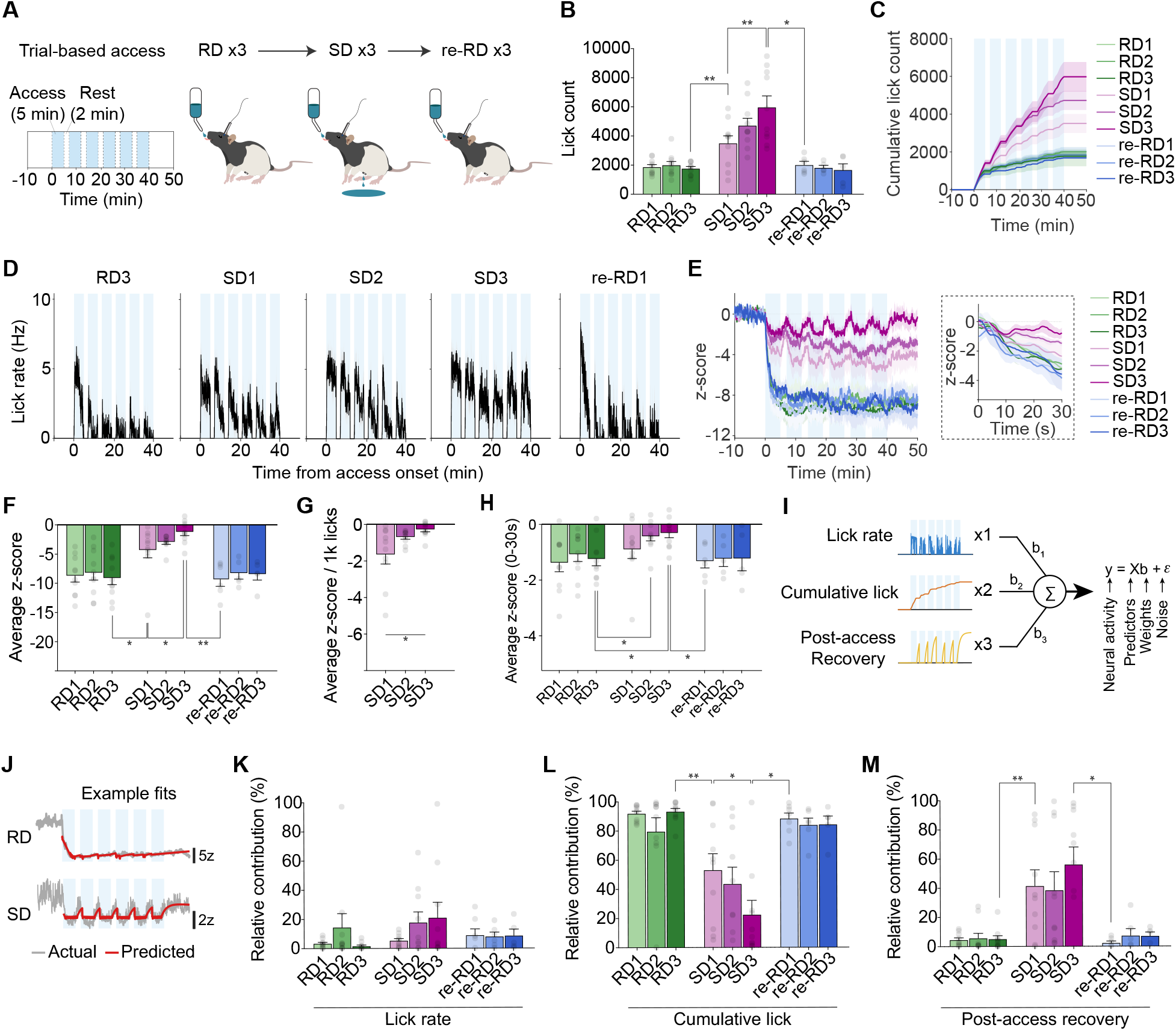
Pre-gastric inhibition of thirst neurons diminishes with learning. **A,** Experimental timeline for repeated trial-based drinking sessions consisting of three real-drinking (RD) sessions, three sham-drinking (SD) sessions, and three return-to-real-drinking (re-RD) sessions. **B,** Total lick count during each RD, SD, and re-RD session. **C,** Cumulative lick count across each RD, SD, and re-RD session. **D,** Average lick-rate traces during repeated access periods in RD3, SD1-SD3, and re-RD1 sessions. **E,** Average MnPO^Agtr1a^ activity traces across repeated RD, SD, and re-RD sessions, with an expanded view of the early access period (0-30 s). **F,** Average MnPO^Agtr1a^ activity in each session. **G,** Average MnPO^Agtr1a^ activity normalized by lick count and expressed per 1,000 (1k) licks across repeated SD sessions. **H,** Average MnPO^Agtr1a^ activity during the first 30 s after access onset across repeated RD, SD, and re-RD sessions. **I,** Regressors included in the generalized linear model (GLM): instantaneous lick rate, cumulative lick count, and post-access recovery. **J,** Representative measured MnPO^Agtr1a^ activity traces overlaid with GLM-predicted traces during RD and SD sessions. **K,** Relative contribution of the instantaneous lick-rate regressor to MnPO^Agtr1a^ activity in each session. **L,** Relative contribution of the cumulative-lick-count regressor to MnPO^Agtr1a^ activity in each session. **M,** Relative contribution of the post-access-recovery regressor to MnPO^Agtr1a^ activity in each session. Sample sizes: RD1-RD3 and SD1-SD3, n = 10 rats; re-RD1, n = 6 rats; re-RD2 and re-RD3, n = 5 rats. \**p* < 0.05, \*\**p* < 0.01. All values are reported as mean ± SEM. More detailed statistical information is in **Table S1**.

We first established a baseline for real drinking, which revealed that thirsty rats consumed most of the water during the first 5 min access period and then licked only sporadically during later periods (**Fig. 5C-D, S4A;** one-way repeated-measures ANOVA: RD1– RD3, all *p* < 0.0001; multiple-comparison test: Access 1 vs. Accesses 2–6, all *p* ≤ 0.025; all comparisons among Accesses 2–6, *p* > 0.05). During the first session of sham drinking, total intake was increased (**Fig. 5B-D, S4E;** lick count: RD3, 1768 ± 142 vs. SD1, 3505 ± 505 licks, *p* = 0.009; consumption: RD3, 13.9 ± 1.7 vs. SD1, 32.5 ± 2.9 mL, *p* = 0.000091), but animals nonetheless showed a reduced rate of drinking in the second half of the session (**Fig. S4D;** Accesses 1–3, 2434 ± 378 licks vs. Accesses 4–6, 1071 ± 171 licks, *p* = 0.0014). This is similar to what we observed for self-paced drinking (**Fig. 1H**) and indicates that some degree of satiation occurred. Total intake increased further during subsequent sham drinking sessions (**Fig. 5B-D, S4E;** lick count: SD1, 3505 ± 505; SD2, 4720 ± 503; SD3, 5974 ± 784; one-way repeated-measures ANOVA, *p* = 0.0061; multiple-comparison test, SD1 vs. SD3, *p* = 0.004) and this was driven primarily by an increase in consumption during the later access periods (**Fig. S4F;** Accesses 1–3: one-way repeated-measures ANOVA, *p* = 0.078; Accesses 4–6: *p* = 0.00033; multiple-comparison test: SD1 vs. SD3, *p* = 0.00045; SD2 vs. SD3, *p* = 0.0026). Finally, when animals were returned to real drinking, the level and pace of consumption immediately returned to the level observed during the initial real drinking session (**Fig. 5B-D, S4C, S4E**; lick count: SD3, 5757 ± 929 vs. re-RD1, 2021 ± 237 licks, *p* = 0.011). Taken together, these data show that the ability of sham drinking to suppress consumption attenuates with experience, but that this is rapidly reversed upon return to real drinking.

We next examined how the activity dynamics of MnPO^Agtr1a^ neurons evolved during these repeated sessions of real and sham drinking. During real drinking, MnPO^Agtr1a^ neurons were rapidly inhibited during the first access period and then remained strongly inhibited for the duration of the session (**Fig. 5E**). This response closely resembled the neural response to self-paced drinking (**Fig. 3E**) and was stable across multiple repeated sessions. When animals engaged in sham drinking for the first time, MnPO^Agtr1a^ neurons were also rapidly inhibited, although the magnitude of this inhibition was reduced compared to real drinking (**Fig. 5E-F**; RD3, −9.1 ± 1.1 z vs. SD1, −4.3 ± 1.3 z; *p* = 0.011). Strikingly, this inhibition progressively diminished across repeated sham drinking sessions (**Fig. 5E-F**; SD1, −4.3 ± 1.3 z; SD2, −3.0 ± 0.4 z; SD3, −1.2 ± 0.6 z; one-way repeated-measures ANOVA, *p* = 0.041; multiple-comparison test: SD1 vs. SD3, *p* = 0.033). This attenuation was already evident during the first 30 s after access onset, with early inhibition weaker across sham drinking sessions than during real drinking (**Fig. 5H**; RD3, −1.24 ± 0.24 z; SD1, −0.90 ± 0.33 z; SD2, −0.43 ± 0.17 z; SD3, −0.31 ± 0.17 z; multiple-comparison test: RD3 vs. SD2, *p* = 0.05; RD3 vs. SD3, *p* = 0.026). This occurred despite the fact that the animals increased their drinking and, as a result, the inhibition per lick sharply decreased across repeated sham-drinking sessions (**Fig. 5G**; SD1, −1.7 ± 0.5; SD2, −0.7 ± 0.1; SD3, −0.3 ± 0.1; one-way repeated-measures ANOVA, *p* = 0.014; multiple-comparison test, SD1 vs. SD3, *p* = 0.013). In contrast, when animals were returned to real drinking, the neural dynamics rapidly reverted to the responses observed in earlier real drinking (**Fig. 5E-F**; SD3, −1.6 ± 1.0 z vs. re-RD1, −9.3 ± 1.2 z; *p* = 0.004). Taken together, these reciprocal behavioral and neural changes indicate that the ability of pre-gastric water cues to inhibit MnPO^Agtr1a^ neurons weakens as rats learn that drinking no longer produces post-gastric rehydration.

We noticed that the dynamics of MnPO^Agtr1a^ neurons during sham drinking had two components: an inhibition that occurred during drinking and a partial recovery that occurred when drinking stopped. To assess the extent to which different features of drinking behavior account for MnPO^Agtr1a^ neural activity, and how this evolves over days, we fit the dynamics to a generalized linear model (GLM) with three regressors: lick rate, cumulative lick count, and post-access recovery, which we defined as an exponential decay from the time of the last lick of each access period (**Fig. 5I-J**). During real drinking, most of the MnPO^Agtr1a^ activity could be explained by cumulative licks alone (**Fig. 5L**; 88.3 ± 3.0% averaged across RD1–RD3; stable across sessions, one-way repeated-measures ANOVA, *p* = 0.24), consistent with the idea that these neurons track total water consumed to produce sustained satiation. During the first session of sham drinking, the contribution of cumulative licks remained substantial (53.1 ± 11.4%) although it was decreased relative to real drinking (**Fig. 5L**; RD3 vs SD1, *p* = 0.007). Upon exposure to subsequent sham drinking sessions, the contribution of cumulative licks further decreased (**Fig. 5L**; SD1, 53.1 ± 11.4%; SD2, 43.6 ± 11.6%; SD3, 22.6 ± 9.8%; one-way repeated-measures ANOVA, *p* = 0.013; multiple-comparison test, SD1 vs SD3, *p* = 0.011). Meanwhile, post-access recovery explained a significant portion of the dynamics during sham but not real drinking (**Fig. 5M**; SD1, 41.5 ± 11.0%; SD2, 38.6 ± 12.6%; SD3, 56.2 ± 11.9%; RD session-averaged, 5.1 ± 1.5%), and this contribution did not change across sessions (**Fig. 5M**; one-way repeated-measures ANOVA, *p* = 0.18). These data are consistent with the idea that loss of post-ingestive feedback attenuates the potency of lick-induced inhibition, which evolves with learning, and simultaneously removes a sustained inhibitory influence that normally suppresses recovery after drinking stops.

### Lick-triggered inhibition of thirst neurons during sham drinking is required for bout termination

We noticed that as consecutive days of sham drinking proceeded, animals required longer bouts of drinking to achieve the same level of neural inhibition (**Fig. 6A**; bout duration × session interaction, linear mixed-effects model, *p* = 0.025; SD1 vs. SD3 slope difference, *p* = 0.019). This was accompanied by a change in behavior in which animals increased drinking by extending the duration of each bout (**Fig. 6B**; SD1, 48.7 ± 4.8 s; SD2, 69.7 ± 8.1 s; SD3, 103.7 ± 19.5 s; one-way repeated-measures ANOVA, *p* = 0.019; multiple-comparison test, SD1 vs. SD3, *p* = 0.015), rather than increasing bout number (**Fig. 6C**; SD1, 16.6 ± 1.5; SD2, 14.5 ± 1.3; SD3, 15.0 ± 2.1 bouts; one-way repeated-measures ANOVA, *p* = 0.60). This suggests that an important role of MnPO^Agtr1a^ neuron inhibition during drinking is to drive the termination of a lick bout, and attenuation of this bout termination signal is a cause of increased drinking.

**Figure 6.**
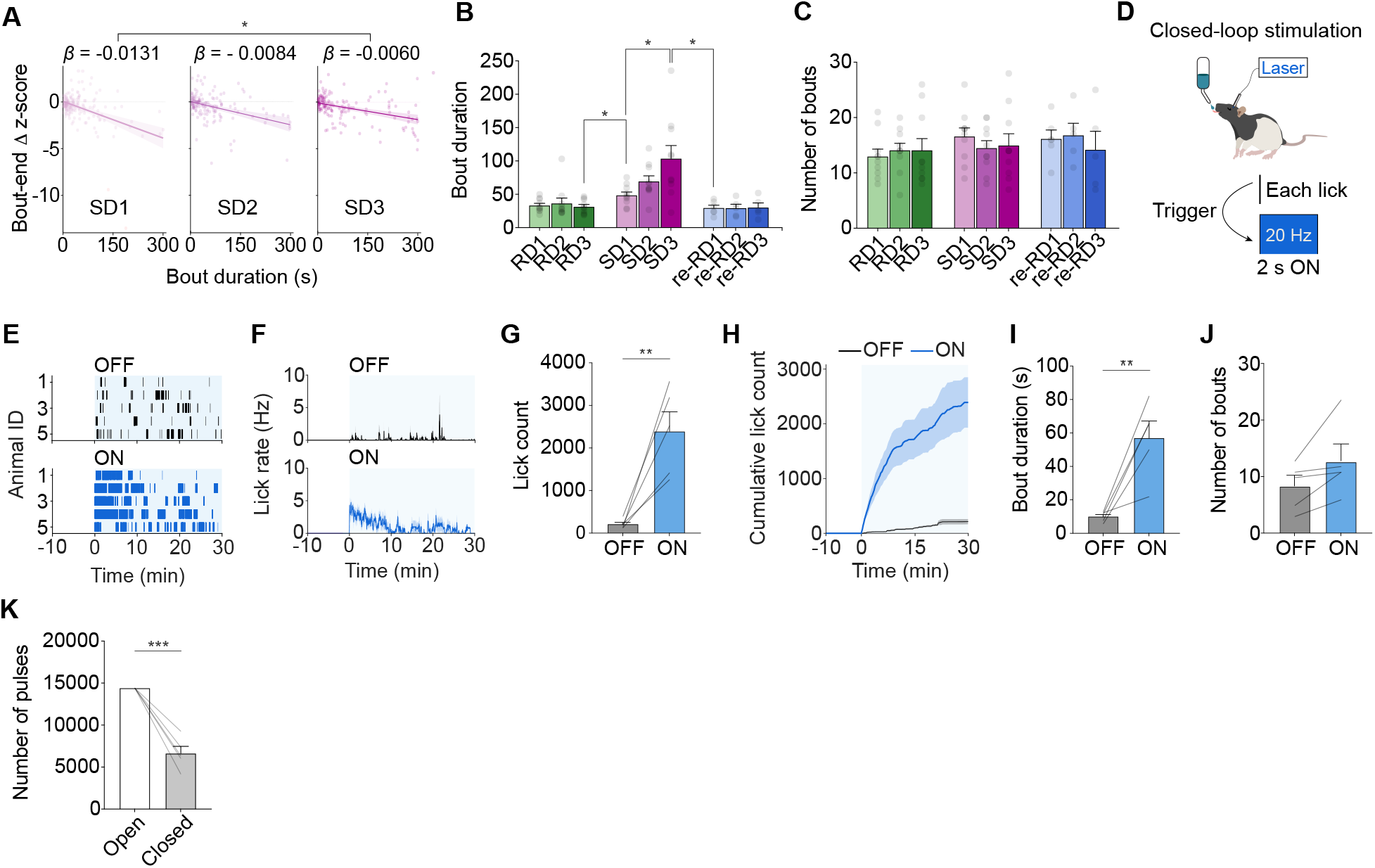
Increased consumption during sham drinking was driven by prolonged drinking bouts. **A,** Relationship between bout duration and the change in MnPO^Agtr1a^ activity at bout end during sham-drinking (SD) sessions SD1-SD3; lines show session-specific linear fits. **B,** Average drinking-bout duration in each real-drinking (RD), SD, and return-to-real-drinking (re-RD) session. **C,** Number of drinking bouts in each session. **D,** Closed-loop optogenetic stimulation protocol in which each lick triggered 2 s of 20-Hz laser stimulation. **E,** Lick rasters from sated rats during 30-min water access in laser-OFF and closed-loop laser-ON sessions. **F,** Lick rate during laser-OFF and closed-loop laser-ON sessions. **G,** Total lick count in laser-OFF and closed-loop laser-ON sessions. **H,** Cumulative lick count in laser-OFF and closed-loop laser-ON sessions. **I,** Average drinking-bout duration in laser-OFF and closed-loop laser-ON sessions. **J,** Number of drinking bouts in laser-OFF and closed-loop laser-ON sessions. **K,** Number of laser pulses delivered during open-loop versus closed-loop stimulation sessions. Sample sizes: **A-C,** RD1-RD3 and SD1-SD3, n = 10 rats; re-RD1, n = 6 rats; re-RD2 and re-RD3, n = 5 rats; **E-K,** n = 5 rats. \**p* < 0.05, \*\**p* < 0.01, \*\*\**p* < 0.001. All values are reported as mean ± SEM. More detailed statistical information is in **Table S1**.

A prediction of this model is that the amount of water consumed should be more strongly driven by MnPO^Agtr1a^ neuron activity during drinking rather than during interbout intervals. To test this idea, we activated MnPO^Agtr1a^ neurons with closed-loop stimulation. Each lick triggered a brief optogenetic stimulation (2 s, 20 Hz; **Fig. 6D**), thereby opposing the normal lick-linked decrease in neural activity. Closed-loop stimulation markedly increased total licking in rats that were not water-deprived (**Fig. 6E-H**; laser OFF, 214 ± 49 licks; laser ON, 2391 ± 462 licks; *p* = 0.0083). This increase was driven primarily by longer drinking bouts (**Fig. 6I**; OFF, 10.1 ± 1.4 s; ON, 57.1 ± 10.1 s; *p* = 0.0099), with little or no consistent change in bout number (**Fig. 6J**; OFF, 8.4 ± 1.9; ON, 12.8 ± 3.0; *p* = 0.08). Notably, closed-loop stimulation promoted twice as much drinking as open-loop stimulation (**Fig. 2G, 6G;** lick count: open-loop, 1187 ± 196 licks; closed-loop, 2391 ± 462 licks; *p* = 0.05), despite delivering fewer than half as many laser pulses (**Fig. 6K**; open-loop, 14,400 pulses; closed-loop, 6,637 ± 836 pulses; *p* = 0.00075). This indicates that MnPO^Agtr1a^ neuron activity is most important during ongoing consumption. Together, these results support a causal role for lick-linked inhibition of MnPO^Agtr1a^ neurons in terminating ongoing drinking bouts and suggest that its attenuation by sham drinking contributes to sustained consumption.

### Sham drinking of hypertonic saline reveals that learning is bidirectional

Repeated sham drinking of water weakens the inhibition of thirst neurons by pre-gastric cues, consistent with the extinction of a learned association between oral water cues and rehydration. We therefore wondered whether learning could proceed in the opposite direction when oral cues predict a post-ingestive consequence that increases thirst. To test this, we allowed thirsty rats to drink a hypertonic solution (0.3 M NaCl) that increases blood osmolality when absorbed (**Fig. 7A**)^22^.

**Figure 7.**
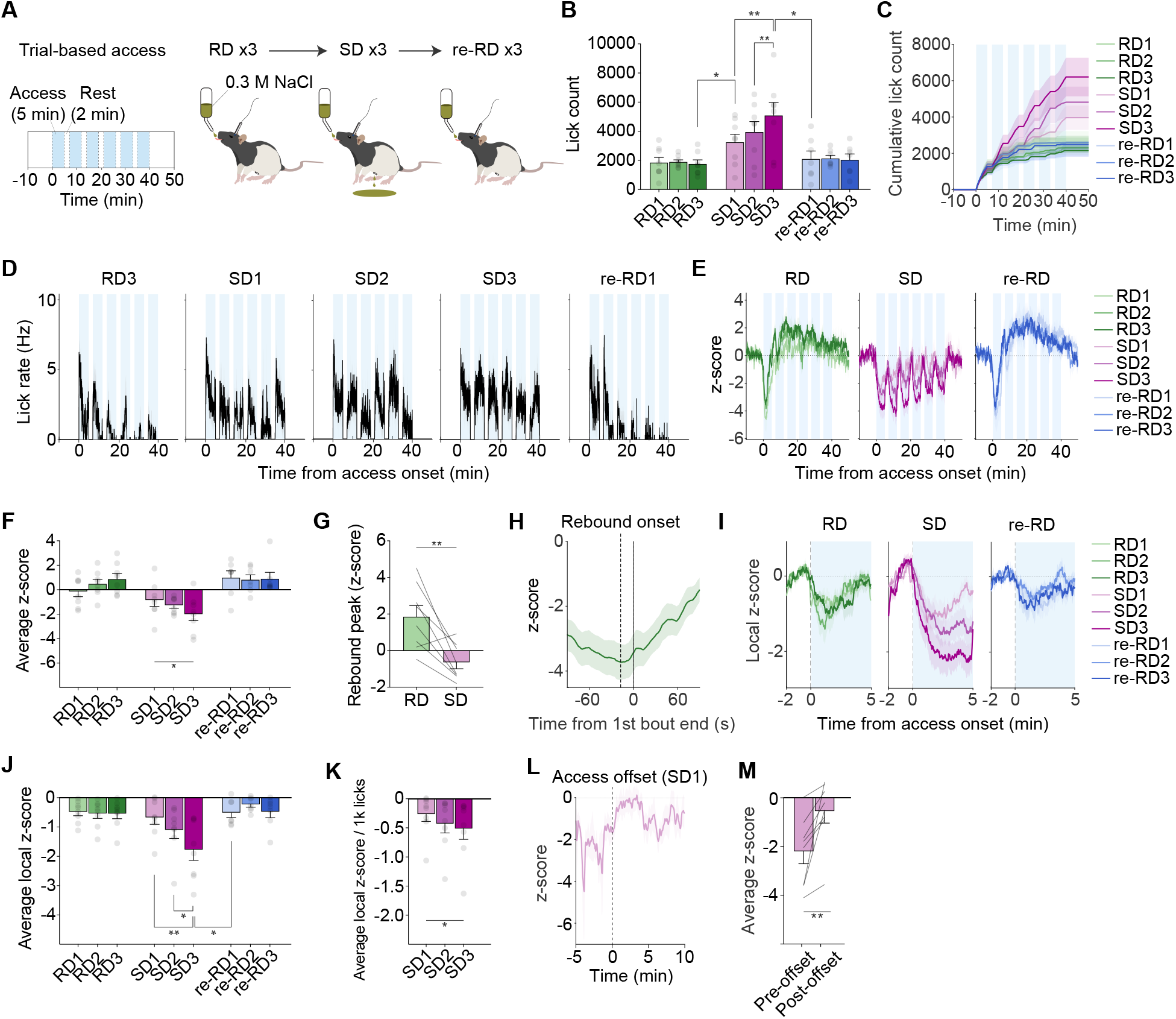
Learning of pre-gastric thirst quenching is bidirectional. **A,** Experimental timeline for repeated trial-based access to 0.3 M NaCl, consisting of three real-drinking (RD) sessions, three sham-drinking (SD) sessions, and three return-to-real-drinking (re-RD) sessions. **B,** Total lick count during each RD, SD, and re-RD session. **C,** Cumulative lick count across each RD, SD, and re-RD session. **D,** Average lick-rate traces during repeated access periods in RD3, SD1-SD3, and re-RD1 sessions. **E,** Average MnPO^Agtr1a^ activity traces across repeated RD, SD, and re-RD sessions. **F,** Average MnPO^Agtr1a^ activity in each session. **G,** Peak MnPO^Agtr1a^ rebound activity during the first access-rest cycle in RD versus SD. **H,** Average MnPO^Agtr1a^ activity trace aligned to the end of the first drinking bout during RD; the vertical marker indicates rebound onset. **I,** Average MnPO^Agtr1a^ activity traces aligned to each access period in RD, SD, and re-RD sessions after local z-scoring to the preceding 2-min baseline. **J,** Average local MnPO^Agtr1a^ activity during each access period across RD, SD, and re-RD sessions. **K,** Average local MnPO^Agtr1a^ activity normalized by lick count and expressed per 1,000 (1k) licks across SD1-SD3. **L,** Average MnPO^Agtr1a^ activity trace in SD1 aligned to the end of the final access period. **M,** Average MnPO^Agtr1a^ activity before and after the final access offset for the trace shown in **(L)**. Sample sizes: RD1-RD3 and SD1-SD3, n = 8 rats; re-RD1-re-RD3, n = 7 rats. \**p* < 0.05, \*\**p* < 0.01. All values are reported as mean ± SEM. More detailed statistical information is in **Table S1**.

Water-deprived rats readily drank the hypertonic solution during real drinking, although there was a trend toward lower consumption of hypertonic saline compared to water (**Fig. 5B–D, S4E, 7B–D, S5A;** 0.3 M NaCl, 9.7 ± 1.7 mL; water, 13.9 ± 1.4 mL; 0.3 M NaCl vs. water, *p* = 0.07). This saline consumption was accompanied by a neural response that had two parts: an initial rapid decline in MnPO^Agtr1a^ activity during drinking, which presumably reflects the satiating effect of pre-gastric cues, followed by a rebound in activity to a higher level than baseline, which may involve post-gastric sensing of the salt content of the fluid (**Fig. 7E**). This rebound occurred with a latency of 86 ± 12 s after drinking onset (RD1, 80.3 ± 15.8 s; RD2, 88.8 ± 20.5 s; RD3, 88.0 ± 11.3 s) and typically preceded the cessation of the first lick bout (**Fig. 7H, S5B;** average across RD1-RD3, 20.8 ± 17.6 s before bout cessation; RD1, 45.2 ± 29.0 s; RD2, 6.3 ± 18.9 s; RD3, 11.0 ± 25.9 s), suggesting that it may contribute to inhibiting further drinking.

We next investigated sham drinking of the same saline solution. Like real drinking, sham drinking was associated with a sharp drop in MnPO^Agtr1a^ activity during initial saline consumption (**Fig. 7E**). However, the response to sham drinking was different in several respects. First, MnPO^Agtr1a^ activity did not rebound until the animals stopped drinking (**Fig. S5C**), suggesting that the rebound during sham drinking reflects release from pre-gastric inhibition as drinking ceases, rather than the emergence of a post-ingestive signal caused by saline absorption. Consistently, in subsequent sham drinking sessions, animals drank almost continuously during the first access period, and, accordingly, the rebound in MnPO^Agtr1a^ activity was delayed until saline access was removed (**Fig. S5C**). Second, whereas the rebound in activity during real drinking exceeded the original baseline (**Fig. 7E, G**; RD rebound peak, 1.86 ± 0.63 z; vs. baseline, *p* = 0.020), during sham drinking activity recovered only to near-baseline levels (**Fig. 7E, G**; SD rebound peak, −0.64 ± 0.35 z; vs. baseline, *p* = 0.106; RD vs. SD, *p* = 0.0059). This is consistent with the fact that only during real drinking does saline consumption result in further bodily dehydration.

We found that the amount of saline consumed increased across consecutive sessions of sham drinking (**Fig. 7B–D, S5A;** consumption: SD1, 28.4 ± 7.7 mL; SD2, 44.4 ± 9.4 mL; SD3, 53.6 ± 9.5 mL; one-way repeated-measures ANOVA, *p =* 0.000028; multiple-comparison test, SD1 vs. SD2, *p* = 0.002; SD1 vs. SD3, *p* = 0.001). However, in contrast to water sham drinking, MnPO^Agtr1a^ inhibition became progressively stronger across sessions, reflecting the extinction of a learned association between oral saline cues and their delayed thirst-promoting hyperosmotic consequence (**Fig. 7I-J**; SD1, −0.68 ± 0.23 z; SD2, −1.10 ± 0.29 z; SD3, −1.78 ± 0.36 z; one-way repeated-measures ANOVA, *p* = 0.00057; multiple-comparison test: SD1 vs. SD3, *p* = 0.001; SD2 vs. SD3, *p* = 0.018). Importantly, this increased inhibition was not solely due to increased drinking, as the inhibition of MnPO^Agtr1a^ neurons per lick also increased with experience (**Fig. 7K**). Moreover, whereas MnPO^Agtr1a^ neurons remained inhibited for at least 15 minutes after sham drinking of water (**Fig. 3K-L**), saline sham drinking was not associated with any durable inhibition of MnPO^Agtr1a^ neuron activity **(Fig. 7L-M**, SD1: pre, −2.22 ± 0.51 z; post, −0.56 ± 0.47 z; pre vs. post, *p* = 0.0091; post vs. baseline, *p* = 0.27). This indicates that the presence of salt blocks the long-lasting, thirst-quenching effect of pre-gastric water cues. Taken together, these data reveal that the satiation of thirst by pre-gastric cues is not a hardwired sensory reflex. Instead, MnPO^Agtr1a^ responses can be flexibly and bidirectionally recalibrated by experience to match the relationship between oral cues and their subsequent physiological consequences.

## Discussion

A central goal of physiology is to explain how hunger and thirst are generated and quenched. Early theories proposed that hunger was the rumbling of the stomach^23^ and thirst was the dryness of the throat^6^. It is now known that these states reflect the global need for energy and water, but the subjective experience of hunger and thirst is nevertheless tightly coupled to such local sensations, and how these somatic cues become linked to general bodily needs is not well understood. A fundamental challenge has been the inability to dissociate these peripheral sensations from broader changes in bodily state during natural ingestion.

Here, we have combined sham drinking, which uncouples water ingestion from rehydration^24–26^, with neural recordings of thirst-promoting neurons in the MnPO. This allowed us to study the connection between peripheral sensations and their central representation during natural drinking behavior. We showed that sham drinking rapidly and strongly inhibits thirst-promoting neurons and terminates consumption despite no water absorption. Remarkably, this neural inhibition by sham drinking tracks cumulative intake and persists for tens of minutes after drinking has stopped, indicating that pre-gastric signals alone are sufficient to estimate the volume of water consumed and translate this into durable inhibition of thirst. We showed that this long-lasting inhibition requires prior passage of water through the oral cavity, as licking or swallowing in the absence of water induced only transient responses. When rats went through repeated experiences of drinking without rehydration, this sustained inhibition of thirst neurons attenuated, revealing that it is learned. Consistently, repeated sham drinking of hypertonic saline, which is dehydrating, had the opposite effect, revealing this learning is bidirectional. Together, these findings reveal that the oropharyngeal detection of water causes long-lasting suppression of thirst, and this flexible suppression reflects a learned relationship between oral cues and post-ingestive consequences rather than a hard-wired sensory response.

### Pre-gastric thirst satiation requires passage of water through the oropharynx

While oropharyngeal cues are clearly important for thirst satiation, the precise nature of these signals has long been debated. The rapid onset of inhibition of thirst circuits by drinking^19,20^ indicates that the signal must originate from the oral cavity^27,28^. It has been proposed that this involves temperature, since oral cooling is thirst quenching^9,29–33^, and taste, since the gustatory system shows water responses^10^. Swallowing (or “gulping”^11^) has been proposed to provide an additional signal that suppresses thirst^28,34–36^, possibly independent of water chemosensation^37^. However, arbitrating between these mechanisms has been challenging due to the difficulty of testing their individual roles in behaving animals. Our findings suggest that this pre-gastric inhibition involves at least two separate signals. The first is a response to the motor act of licking, independent of fluid ingestion, which can weakly and transiently inhibit thirst neurons during the first minutes of licking (**Fig. 4B-D**). The second, and more important, signal is tightly linked to the passage of water through the oral cavity and esophagus. This signal is responsible for most of the rapid inhibition in the first minutes of drinking and all the long-lasting inhibition that persists after drinking stops. Importantly, this signal requires the chemosensation of water, because ingestion of corn oil, which triggers vigorous licking and swallowing, nevertheless produces no additional thirst neuron inhibition beyond licking an empty sipper. This indicates that swallowing is insufficient for thirst quenching and that the chemical detection of water is critical^10^.

Oropharyngeal cues function in concert with gastrointestinal signals to mediate the rapid quenching of thirst. Previously, we and others showed that these gastrointestinal signals emerge with a delay of approximately one minute from the onset of drinking and track the osmolarity of ingested fluid^11–13^. Together, these observations led to an intuitive “sequential model” in which oropharyngeal and gastrointestinal signals cooperate to drive thirst quenching on different timescales^38^: oral cues act immediately, but transiently, to inhibit thirst neurons during the act of drinking, whereas subsequent gastrointestinal signals “stabilize” this rapid inhibition and allow it to persist for minutes until the ingested water is absorbed.

Here, we have tested this model by measuring for the first time how oral water ingestion affects thirst neuron activity in the absence of subsequent gastrointestinal feedback. Surprisingly, we found that oropharyngeal cues alone can cause strong inhibition of thirst neurons that lasts for tens of minutes after drinking has stopped. This indicates that the sequential model is incorrect, and that early oral cues are sufficient to produce long-lasting neural responses. Nevertheless, the fact that sham drinking results in weaker thirst neuron inhibition than real drinking, and causes corresponding over-drinking, confirms that gastrointestinal feedback is important. In the future, it will be important to establish methods that allow, similar to sham drinking, the clean separation of gastrointestinal and post-absorptive signals during natural water ingestion, in order to clearly define their respective contributions.

### The rapid satiation of thirst by oral cues is learned, not hardwired

The feeling of ingested water rapidly quenching thirst is a fundamental and universal aspect of human experience. Moreover, thirsty animals show appetitive responses to oral water from the earliest ages that can be tested, and this develops before motivated responses to food^39,40^. Thus, it is intuitive to assume that rapid thirst satiation is an innate response to ingestion of water. However, there is also evidence that animals can learn about the rehydrating properties of different fluids^25,41^. This raises the question to what extent the fundamental sensation of oral thirst quenching is a hardwired reflex versus a flexible, plastic response.

We found that, across repeated sessions of sham drinking, the ability of water to inhibit MnPO^Agtr1a^ neurons declined with experience. This manifested as a decrease in the neural inhibition per lick of water, indicating the satiating power of water itself diminished over time. As this learning occurred, animals required more licks to achieve the same degree of neural inhibition per bout, and, accordingly, the length of each bout progressively increased. In parallel, we showed that repeated sham drinking of hypertonic saline, which allowed animals to learn that this fluid is not dehydrating as expected, resulted in progressively stronger MnPO^Agtr1a^ inhibition. These data indicate that the value of oral fluid cues can be learned based on post-ingestive consequences. This learning is bidirectional and reflected in changes in the lick-triggered dynamics of thirst-promoting neurons in MnPO.

An initially surprising feature of the saline experiments was that saline intake increased across sham-drinking sessions even as MnPO^Agtr1a^ inhibition became stronger. However, these changes are not necessarily contradictory because sham drinking alters the relationship between oral cues and post-ingestive consequences differently for saline than for water. During real saline drinking, oral saline cues are followed by a delayed hyperosmotic consequence that activates MnPO^Agtr1a^ neurons and limits further intake. Repeated sham drinking removes this adverse consequence, allowing animals to learn that saline consumption no longer predicts a worsening of their physiological state and thereby promoting greater saline intake. At the same time, the stronger MnPO^Agtr1a^ inhibition could reflect weakening of a learned excitatory component normally associated with the delayed hyperosmotic consequence of saline, strengthening of the inhibitory influence of saline-associated sensory signals, or both. Our experiments do not distinguish between these possibilities. Nevertheless, the parallel changes in behavior and MnPO^Agtr1a^ activity support a model in which learning updates the predictive meaning of oral fluid cues according to their later physiological consequences, rather than simply scaling thirst-neuron inhibition in proportion to how much an animal ultimately drinks.

These findings establish an important role for learning in thirst satiation, but they do not establish how this learning happens mechanistically. One important question is which post-ingestive effect of water ingestion functions as the unconditioned stimulus. This could involve sensing of ingested water in the intestine or hepatic portal circulation, which is known to contribute to thirst satiation^11–13^, and/or water absorption into the blood, which is associated with delayed dopamine release and modulates fluid preferences^41–43^. A second question is where in the brain the learning happens and which plasticity mechanisms are involved. We have previously shown that individual MnPO neurons receive separate signals reflecting detection of water in the mouth, GI tract, and blood^12^. This convergence of pre-gastric and post-ingestive cues raises the possibility that the MnPO could be an important site of plasticity. On the other hand, it is possible this occurs at upstream nodes in the thirst circuit in the lamina terminalis, pons or medulla^44^. Answering this question will require investigating these upstream nodes, including their necessity for learning, dynamics during sham drinking, and synaptic properties.

## Materials and Methods

All experimental protocols were approved by the University of California, San Francisco Institutional Animal Care and Use Committee (IACUC) following the National Institutes of Health guidelines for the Care and Use of Laboratory Animals.

### Rat strains

Male and female Long-Evans rats (>10 weeks old) were used in all experiments. Experiments were performed between 8:00 am and 4:00 pm. Standard lab chow and water were provided ad libitum in home cages. Experimental animals were singly housed under a 12-h light/12-h dark cycle (6:00-18:00) at constant room temperature. Breeding animals can be either singly housed or group housed in the animal facility.

We obtained both male and female wild-type Long-Evans control rats (Charles River Laboratories Strain Code: 006L/E), tdTomato reporter rats (Dr. Hiromu Yawo at Tohoku University, Strain name: LE-Tg(Gt(ROSA)26Sor-CAG-tdTomato)24Jfhy).

### Genotyping and Identification

Tissue for genotyping was collected either from the ear using a commercial ear punch (2mm diameter) or by tail snipping with a small pair of scissors. These ear punches also allowed for the identification of animals from the same litter.

### Generation of Agtr1a^Cre^ rats

Agtr1a^Cre^ rats were generated by homologous recombination at the endogenous Agtr1a genomic locus mediated by targeted CRISPR endonuclease activity. Homology regions were captured into a plasmid from a BAC containing the Agtr1a locus by recombineering. The T2A-Cre sequence was inserted immediately upstream of the endogenous stop codon. The final targeting vector containing the 5’ and 3’ homology arms was verified by restriction digest and sequencing. To generate site-specific double-stranded breaks using CRISPR, an sgRNA was selected such that the guide sequence would be separated from the PAM site in the genomic DNA by the T2A-Cre insertion. This ensured that the targeting vector and recombined Agtr1a-T2A-Cre allele were protected from Cas9 nuclease activity. Super-ovulated female rats were mated to stud males, and fertilized zygotes were collected from the oviducts. Cas9 protein, sgRNA, and targeting vector DNA were mixed and injected into the pronucleus of fertilized zygotes. Injected zygotes were implanted into the oviducts of pseudopregnant female rats. Pups genotyped positive for the knock-in allele were kept as founders. Two independent knock-in lines were crossed to the reporter line, and reporter expression patterns from these lines were identical. All Agtr1a-T2A-Cre rats used here were maintained on a Long-Evans background. Founder pups and offspring were genotyped for the presence of the knock-in allele by qPCR. Presence of the knock-in allele was also verified by long-range PCR.

### Gastric fistula implant surgery

The day before surgery, rats were food-restricted overnight for about half of their regular consumption. Water was still given ad libitum. The stainless-steel cannula, screwcap, and hex nut, as well as all surgical equipment, were autoclaved. Rats were anesthetized with isoflurane and room air (induction 5%; maintenance 1.8-3%) using the Somnosuite (Kent Scientific Item # SS-01). Ophthalmic ointment was applied to the eyes. Subcutaneous injections of Carprofen (5 mg/kg), Enrofloxacin (8 mg/kg), and Atropine (0.05 mg/kg) were given. The body temperature was maintained with a heating pad underneath the body and monitored with a rectal thermal sensor. Rats were placed in the prep area with their abdomen facing up. The ventral surface of the rat was shaved, and then Nair Hair Remover was applied to remove the remnant hair. The animals were then transferred to the surgery station, and the skin was subsequently scrubbed with betadine and alcohol three times. Following adequate depth of anesthesia (as monitored via the toe pinch reflex), 100 μL of 1% lidocaine was injected subcutaneously about 1 inch below the xiphoid process. After about five minutes, a midline incision with a #10 scalpel blade was made through the skin and the linea alba of the abdominal muscles. The lower part of the stomach was grasped with a blunt forcep to pull out the stomach. The stomach was then placed on a sterile gauze pad soaked with sterile saline to prevent it from sliding back under the liver. For proper drainage of stomach contents, the cannula was placed at the point where the limiting ridge meets the greater curvature of the stomach. The proximal part of the stomach just to the right (visual right) of the limiting ridge was grasped, and a small (∼2 mm) hole was cut through the wall of the stomach with an iris scissor. While holding the left edge of the incision with a blunt forceps, the circular base of the cannula was inserted with a clockwise twist using gentle pressure. When the base of the cannula is completely inside the stomach, gauze or a cotton swab is inserted into the lumen of the cannula to prevent leakage of gastric contents. If leakage occurred, we would flush the peritoneal cavity immediately with sterile isotonic saline to prevent peritoneal irritation by gastric acid. A purse-string suture (4-0 monofilament sutures) was placed through the wall of the stomach around the base of the cannula ∼5 mm from its shaft. The free ends of the suture were tied, and the ends of the stomach wall were inverted into the stomach so that the outer (serosal) surface of the stomach fits tightly against the shaft of the cannula. A small incision (∼2 mm) was made with a #10 scalpel blade about 2 inches below the armpit through the skin and abdominal wall. A 1 mL pipette tip was attached to the pointed end of the cannula, then the cannula was guided through the incision to penetrate the abdominal wall and the skin until resistance occurred due to the base of the cannula pressing against the peritoneal surface of the abdominal muscles. Saline can be applied to the pipette tip to reduce friction. Once the cannula is exteriorized, the pipette tip is detached, and the screw cap is inserted into the lumen of the cannula to close it. The area of skin around the base of the cannula shaft will be coated with triple antibiotic ointment. A hex nut was placed outside the shaft of the cannula and threaded downwards until it touched the skin to prevent the cannula from sinking back. The incision was closed through the muscle layer with 3-0 monofilament sutures. The skin was closed with wound clips. During surgeries, the rat’s body temperature, breathing, and toe pinch reflexes were continuously monitored. Sustained-release buprenorphine (Ethiqa XR, 0.65 mg/kg) was given to each rat at the beginning of the surgery.

### Sham drinking behavior

The rats were water-restricted 24 h prior to behavioral sessions. The cannula was unscrewed, and the stomach was flushed with lukewarm 0.9% saline (prepared from salt and double-distilled water) (37 °C) by inserting a 20-mL syringe into the lumen of the cannula. The saline was injected quickly and with slight pressure. We flushed until no food particles were in the gastric drainage after two consecutive injections of saline. A drainage adapter was attached by threading it into the cannula. The rats were then gently placed into a behavioral testing chamber (Med Associates, ENV-007CT). The weight of the gastric drainage was measured as well as the weight of the liquid consumed. After a 10-min baseline period, a licking spout connected to a bottle was delivered via a retractable lickometer (Med Associates, ENV-252M). A typical test session lasted 40 minutes to 2 hours. At the end of the test, the screwcap was replaced, and the rat returned to its home cage. Maintenance chow and water were restored. Animals were given a minimum of 48 hours between tests.

We used three measures to evaluate the completeness of gastric drainage during SD of water: (1) Gastric drainage of ingested liquid must begin within the first 15 s of ingestion; (2) The total volume of gastric drainage must equal or exceed 85% of the volume ingested; (3) When the drainage adapter is removed from the cannula at the end of a test, a significant volume of drainage must not occur. When these criteria were satisfied, gastric drainage was considered complete or nearly complete. If all three criteria were not met, drainage was considered incomplete, and the results of that test were discarded. Incomplete drainage was rare and if this occurred repeatedly the rat was then removed from the experiment.

### Viruses

The following viruses were purchased from Addgene: AAV1-EF1a-double floxed-hChR2(H134R)-mCherry-WPRE-HGHpA (Plasmid #20297), AAV1-hSyn-DIO-mCherry (Plasmid #50459), AAV1-CAG-Flex-GCaMP6s-WPRE-SV40 (Plasmid #100842). All viruses were diluted with sterile PBS to ∼1-5 × 10^12^ vg/mL before injection. Injection volume was 250-300 nL. Animals were allowed 4-6 weeks after stereotaxic injection of the viruses for recovery and viral expression. The injection coordinates for MnPO were AP −0.12 mm, ML 0, from the bregma, and DV −6.4 mm from the surface of the skull.

### Intracranial surgery

All surgical equipment was autoclaved. Rats were anesthetized with isoflurane and room air (induction 5%; maintenance 1.8-3%) using the Somnosuite (Kent Scientific Item # SS-01). Ophthalmic ointment was applied to the eyes. Subcutaneous injections of Carprofen (5 mg/kg), Enrofloxacin (8 mg/kg), and Atropine (0.05 mg/kg) were given. The body temperature was maintained with a heating pad underneath the body and monitored with a rectal thermal sensor. The scalp was shaved and then scrubbed with betadine and alcohol wipes three times. The animal was then transferred to the stereotaxic head frame (KOPF Model 940). 100 µL of 1% lidocaine was injected subcutaneously along the midline before an incision was made with a #10 scalpel blade. Craniotomy was made with a dental drill (0.6 mm drill bit). Virus was injected with a Micro4 microsyringe pump controller (WPI, Sarasota, FL) using a 10 µL Hamilton syringe (WPI, Sarasota, FL) and a glass pipette at the rate of 125 nL/min. The glass pipette was held at the injection site for 10 min after the injection was finished before being retracted slowly. If no implant was needed after viral injection, the scalp was closed with 4-0 monofilament sutures.

### Fiber photometry

Following the craniotomy and viral injection, the glass pipette was held at the injection site for 10 min after the injection was finished before being retracted slowly. Four additional craniotomies were made on four different skull plates. Screws were inserted into the craniotomy opening. A 400 µm diameter mono fiber optic cannula (Doric Lenses, MFC_400/430-0.48_8mm_MF2.5_FLT) was implanted 200 µm above the viral injection site. Then Metabond dental cement (Patterson Dental Supply INC: 07-5533559, 07-5533500; Henry Schein Inc.: 1864477) was applied to cover the base of the optic cannula and four screws. Lastly, flow-It (Patterson Dental Supply INC: 07-6472542) was applied and cured with a UV lamp as an additional layer of fortification and protection for the implant.

During fiber photometry recordings, a low-autofluorescence patchcord (Doric Lenses, MFP_400/430/ARMO-0.57_3m_FCM-FCM_LAF) was tethered to the implanted cannula via an adapter (Doric Lenses, ADAPTER_FC_RO). Continuous 6 mW blue LED (470 nm) and UV LED (415 nm) served as excitation light sources. These LEDs were driven by a multichannel hub (Thorlabs), modulated at 211 Hz and 511 Hz, respectively, and delivered to a filtered minicube (Doric Lenses, FMC6_AE(400-410)_E1(450-490)_F1(500-540)_E2(550-580)_F2(600-680)_S) before connecting through optic fibers. GCaMP6s signals and UV isobestic signals were collected through these same fibers into a femtowatt silicon photoreceiver (Newport, 2151). Digital signals were sampled at 1.0173 kHz, demodulated, lock-in amplified, sent through a processor (RZ10X, Tucker-Davis Technologies), and collected by the Synapse software (TDT).

### Optogenetics

Following the craniotomy and viral injection, the glass pipette was held at the injection site for 10 min after the injection was finished before being retracted slowly. Four additional craniotomies were made on four different skull plates. Screws were inserted into the craniotomy opening. A 400 µm diameter mono fiber optic cannula (Doric Lenses, MFC_400/430-0.48_8mm_SM3_FLT) was implanted 400 µm above the viral injection site. Then Metabond dental cement (Patterson Dental Supply INC: 07-5533559, 07-5533500; Henry Schein Inc.: 1864477) was applied to cover the base of the optic cannula and four screws. Lastly, flow-It (Patterson Dental Supply INC: 07-6472542) was applied and cured with a UV lamp as an additional layer of fortification and protection for the implant.

During optogenetic stimulation, a low-autofluorescence patchcord (Doric Lenses, MFP_400/430/ARMO-0.57_3m_FCM-CM3_LAF) was screwed onto the implanted cannula. For open-loop stimulation and pre-stimulation experiments, the laser was modulated at 20 Hz for a 2 s ON and 3 s OFF cycle with a 10 ms pulse width. For closed-loop stimulation, the laser was turned on for 2 s after each lick was detected by the Med Associates software (MED-PC V) at 20 Hz with a 10 ms pulse width. Laser was delivered by a DPSS 473-nm laser (Shanghai Laser and Optics Century BL473-100FC) through a mono fiber optic patch cord at a laser power of 10 mW, which was measured by a power meter and sensor (Thorlabs, PM100D and S130C) at the tip of the patch cord before the beginning of each experiment each day.

### Lever-pressing

For the negative-reinforcement experiment, sated rats were stimulated with a laser constantly at 20 Hz with a 10 ms pulse width. Two identical-looking retractable levers (Med Associates, ENV-112CM) were placed inside the behavior chamber. After a 10 min baseline period, the levers were extended to allow animals to press. Pressing the active lever turned off the laser for 1 min, while pressing the inactive lever had no effect. Total length of the session was 4 h. The number of presses from each lever was recorded by the Med Associates software (MED-PC V).

### Real-Time Place Preference

A two-chamber apparatus was customized from a rat cage (Ancare, R20PC, 10½” W x 19” L x 8” H). The two sides differed in wall patterns (dot versus stripe pattern). A cardboard divider was placed in the middle of the cage with an opening to allow free access to either side of the apparatus. The videos were taken by a webcam hanging above the cage. Each session was recorded for 30 min. The side that was paired with laser stimulation was counter-balanced between animals. Laser stimulation was at 20 Hz with a 10 ms pulse width when the animal resided on the paired side. A custom MATLAB script was used to track the center-of-mass of the rats to calculate the percentage of time spent on each side of the apparatus.

### Histology

Rats were anesthetized with isoflurane and then transcardially perfused with phosphate-buffered saline (PBS) and 4% formalin. The carcass was then decapitated. The head was stored in 4% formalin overnight before the brain was dissected. Then the brain was stored in 4% formalin again overnight before being moved to 30% sucrose in PBS. After at least 48 h, brains were sectioned at a thickness of 50 μm using a cryostat (CryoStar NX70) and mounted onto slides with DAPI Fluoromount-G (Southern Biotech). Viral expression and lens placement were then evaluated using either a wide-field microscope (Nikon Ti2) or a confocal microscope (Zeiss LSM 710).

### Plasma osmolality measurements

Rats were physically restrained inside a shoe cover. The tip of the tail was snipped with scissors 15 minutes after a real drinking, sham drinking, or a mock (sated or water-deprived) session. 200 µL of blood was collected into EDTA-coated capillary tubes (MICROVETTE 200 EDTA PK100). The blood collection process took on average 1 min. Manual pressure was applied with a gauze sponge to stop any bleeding afterward. The collected blood was immediately put on ice and then centrifuged at 2,000 x g for 10 min at 4 °C. Around 50 µl of plasma was carefully extracted from the top layer with a 100 µL pipette to avoid disturbing the bottom layer of blood cells. The plasma was kept on ice and immediately measured in triplicate using a Vapor Pressure Osmometer (VAPRO® 5600).

### X-ray fluoroscopy

X-ray fluoroscopy recordings were done using a low-energy fluoroscope (Glenbrook Technologies, Newark, NJ). Fluoroscopy data were acquired at 30 frames/s with a voltage of 25 kV and a current of 0.216 mA. Rats were habituated to a custom-made box with a drinking bottle. Water or Ensure was mixed with the oral contrast barium 980 suspension (barium sulfate oral suspension 60% w/v, Liquid E-Z-PAQUE, Bracco Diagnostics) for visualization under X-ray fluoroscopy.

### Fiber photometry data analysis

Fiber-photometry data collected in TDT Synapse software were analyzed in MATLAB using custom scripts. GCaMP and isosbestic signals were downsampled before analysis, and the first 10 min of each recording were excluded as habituation. To correct for shared fluctuations, including movement artifacts and photobleaching, the isosbestic signal was fit to the GCaMP signal by robust linear regression using the 10-min pre-access baseline. Fluorescence was then expressed as:

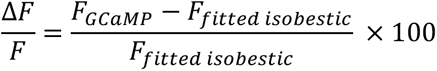

The resulting ΔF/F signal was converted to a z-score using the mean and standard deviation of the same 10-min pre-access baseline. Lick timestamps and water-access periods were synchronized with the photometry signal.

### Generalized Linear Model (GLM)

To determine how drinking-related variables accounted for MnPO^Agtr1a^ activity, each session was analyzed using a ridge-regularized GLM with three regressor families. Lick rate represented ongoing licking and was calculated in 1-s bins. Cumulative lick count represented the running number of licks across the session. Post-access recovery represented the gradual return of neural activity after the final lick of each access period and was modeled using an exponential recovery function beginning at the time of the final lick. The recovery regressor was scaled only during its active post-lick interval, reset at the onset of the next access period, and constrained to zero otherwise. Other regressors were standardized within each session. Models were fit to the z-scored photometry signal over the full post-baseline period using ridge regression with a fixed regularization parameter of λ = 10³. Three nested models were evaluated: lick rate alone; lick rate plus cumulative lick count; and a full model containing lick rate, cumulative lick count, and recovery. Model performance was quantified using adjusted R^2^. The unique contribution of each regressor family was estimated by removing that family from the full model and calculating the resulting reduction in adjusted R^2^:

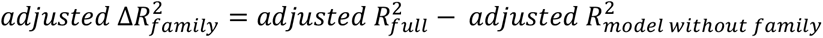

Relative contributions were calculated by dividing each family’s positive adjusted ΔR^2^ by the sum of positive adjusted ΔR^2^ values across the three regressor families. Model fits and regressor contributions were calculated separately for each animal and session before group-level statistical analysis.

## Statistics

Detailed statistical information for each figure is summarized in **Table S1**, complementing the statistical results reported in the main text. No formal statistical methods were used to predetermine sample sizes; however, sample sizes were consistent with those commonly used in the field. For parametric tests, data were assumed to follow a normal distribution given the robustness of these tests to moderate deviations from normality. Statistical tests were two-sided unless otherwise specified.

For within-subject pairwise comparisons, two-tailed paired Student’s *t*-tests were used, whereas unpaired Student’s *t*-tests were used for comparisons between independent groups. One-sample *t*-tests were used to compare values with a defined baseline. For analyses involving two repeated-measures factors, two-way repeated-measures ANOVA was used. For analyses involving one repeated-measures factor, one-way repeated-measures ANOVA was used, whereas one-way ANOVA was used for comparisons involving one non-repeated factor. Post hoc multiple-comparison tests were performed as appropriate. Pearson’s correlation was used to assess linear relationships between continuous variables. Exact statistical tests, sample sizes, test statistics, and *p* values are provided in **Table S1**.

## Acknowledgements

We thank Dr. Yada Treesukosol for instruction in gastric fistula surgery and for sharing component designs. We thank Drs. Joshua Berke, Vijay Mohan Namboodiri, and Evan Feinberg for feedback throughout the course of this study. We also thank Drs. Brooke Jarvie, Christopher Zimmerman, and James Grove for their constructive comments on the manuscript. This work was supported by R01-DK106399, R01-DK145100, and R01-DK138127 (Z.A.K.) and by an AAN Neuroscience Research Training Scholarship (H.N.). Z.A.K. is an Investigator of the Howard Hughes Medical Institute.

## Author Contributions

Z.L. and Z.A.K. conceived the project. Z.L., H.N., and Z.A.K. designed experiments. Z.L. and J.Z. led and performed experiments. Z.L., H.N., and Z.A.K. analyzed and interpreted data. H.N. performed all formal data analyses and generated all figures. Z.L. and Z.A.K. generated and validated the Agtr1a-2A-Cre rat. Z.L., J.Z., S.L., and J.D. managed the rat colony and genotyping. Z.L. performed all stereotaxic surgeries. Z.L., J.Z., and N.L.S.M. performed gastric fistula implant surgeries. Z.L., J.Z., and X.W. performed photometry, optogenetics, and behavioral experiments. Z.L., J.Z., and T.C. performed plasma osmolality measurements. Z.L., J.Z., and A.C. collected and analyzed histology. Z.L., H.N., and Z.A.K. wrote the manuscript with input from all authors.

## Competing financial interests

The authors declare no competing interests.

## Additional information

Correspondence and requests for materials should be addressed to Zachary A. Knight.

## Data availability

The data from this study are available from the corresponding author on reasonable request.

## Code availability

Links to the code used for data analysis are found at https://github.com/zackknightlabucsf/zackknightlabucsf.

---

**Figure S1 (related to Fig. 1).**
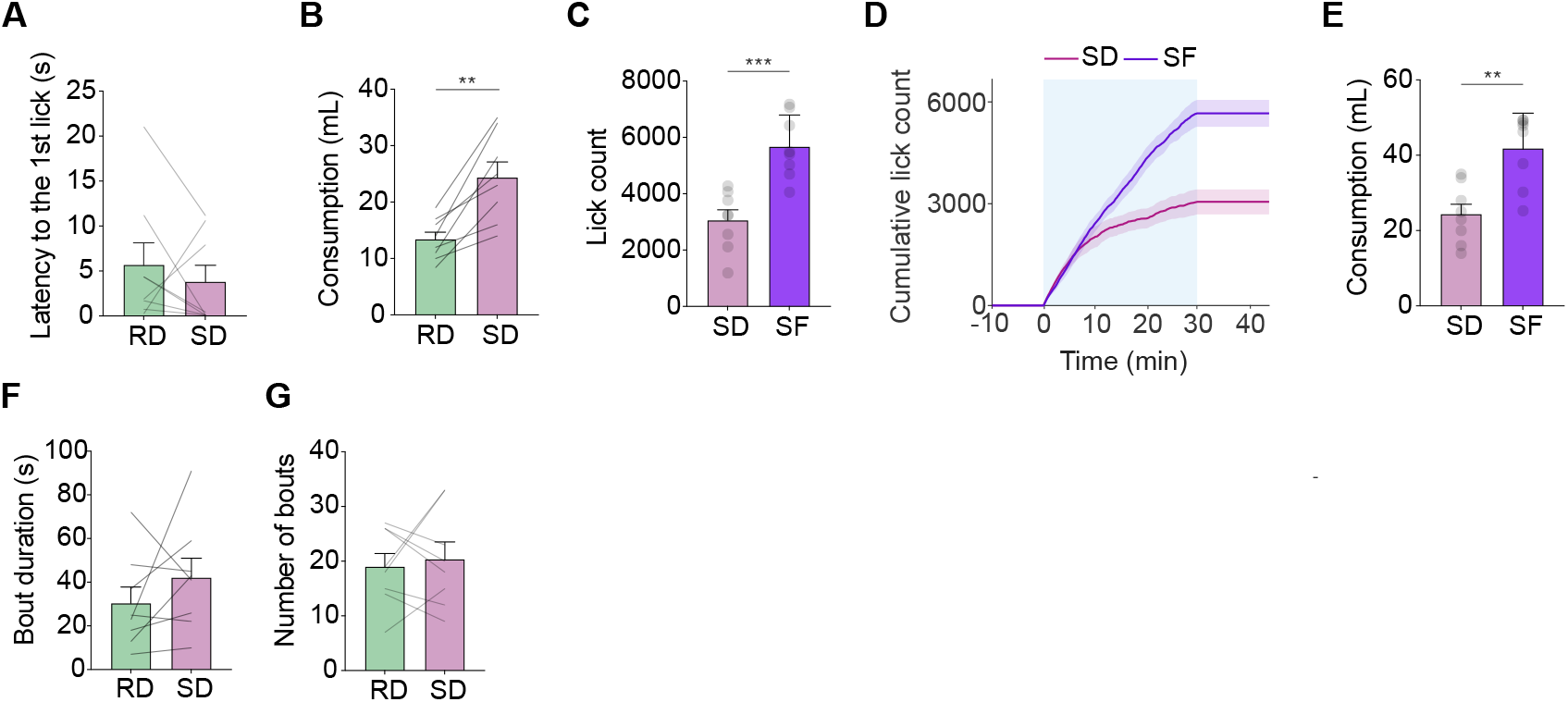
Comparisons between consumption during real drinking, sham drinking, and sham feeding of sucrose. **A,** Latency from water access onset to the first lick during real drinking (RD) versus sham drinking (SD). **B,** Water consumption during RD versus SD. **C,** Total lick count during SD versus sham feeding (SF) of 0.8 M sucrose. **D,** Cumulative lick count during SD and SF. **E,** Fluid consumption during SD and SF. **F,** Average bout duration during RD and SD. **G,** Number of bouts during RD and SD. Sample sizes: n = 8 rats. \*\**p* < 0.01, \*\*\**p* < 0.001. All values are reported as mean ± SEM. More detailed statistical information is in **Table S1**.

**Figure S2 (related to Fig. 2).**
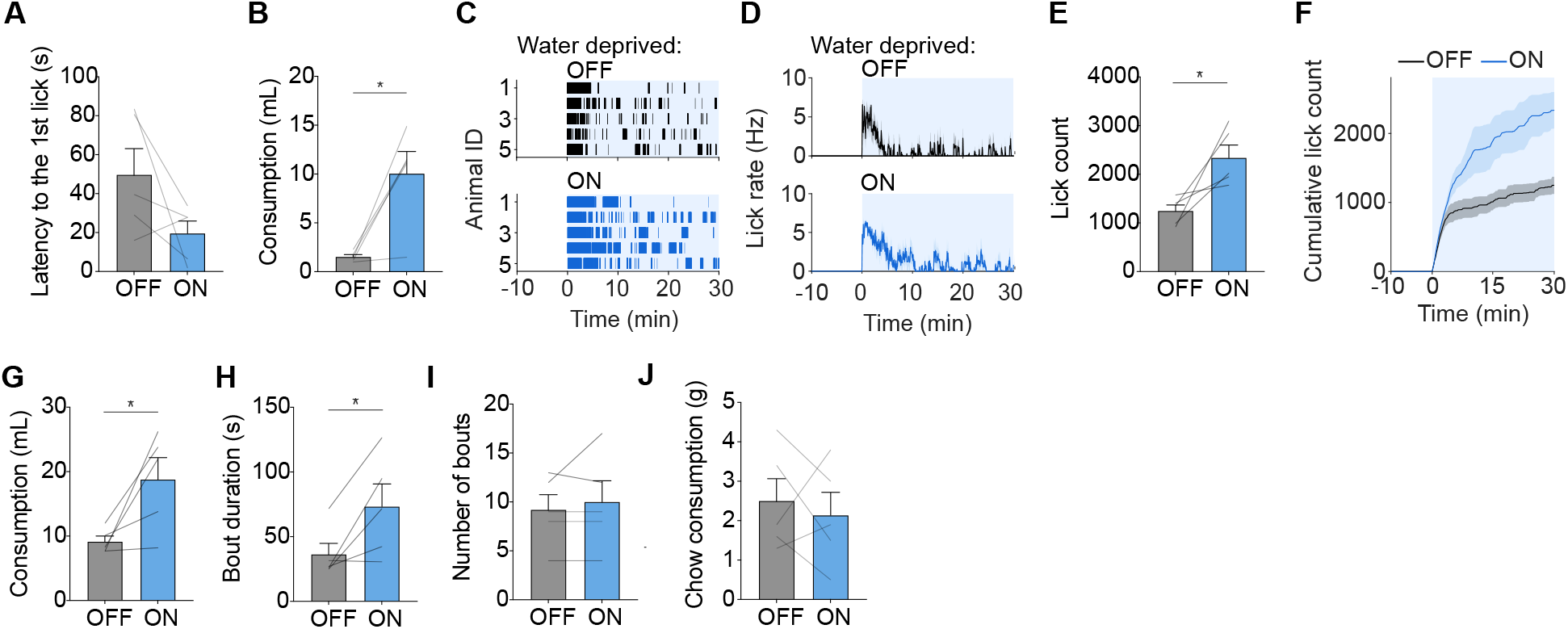
Additional optogenetic functional validation experiments. **A,** Latency from water access onset to the first lick during laser-OFF versus laser-ON sessions. **B,** Water consumption by sated rats during laser-OFF versus laser-ON sessions. **C,** Lick rasters from water-deprived rats during 30-min water access in laser-OFF and laser-ON sessions. **D,** Lick rate during 30-min sessions in water-deprived rats during laser-OFF and open-loop laser-ON sessions. **E,** Total lick count in each session. **F,** Cumulative lick count in each session. **G,** Water consumption by water-deprived rats during laser-OFF versus laser-ON sessions. **H,** Average bout duration in each session. **I,** Number of bouts in each session. **J,** Chow consumption during 30-min sessions in hungry rats during laser-OFF and open-loop laser-ON sessions. Sample size: n = 5 rats. \**p* < 0.05. All values are reported as mean ± SEM. More detailed statistical information is in **Table S1**.

**Figure S3 (related to Fig. 5).**
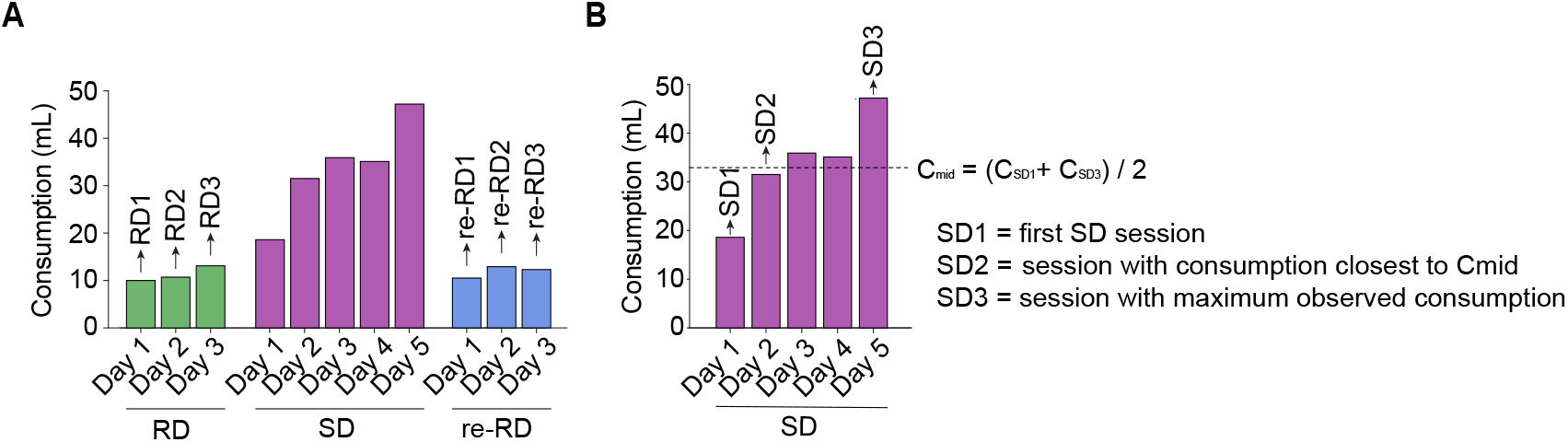
Selection of three sessions for repeated RD, SD, and re-RD analyses in the water-drinking paradigm. **A,** Consumption across consecutive real drinking (RD), sham drinking (SD), and re-RD sessions from a representative animal. Three RD sessions were performed before SD, and three re-RD sessions were performed after SD; all RD and re-RD sessions were included in subsequent analyses. **B,** Behavioral alignment of SD sessions. Because the rate of behavioral adaptation during repeated SD varied across animals, three SD sessions were selected based on consumption to represent comparable early, intermediate, and late stages of adaptation, rather than using fixed chronological sessions. SD1 was defined as the first SD session and SD3 as the session with maximum observed consumption. The midpoint consumption was calculated as C_mid_ = (C_SD1_+ C_SD3_) / 2, and SD2 was defined as the intervening session with consumption closest to C_mid_. This procedure yielded three behaviorally aligned stages representing early, intermediate, and late SD adaptation for comparison with the three selected RD and re-RD sessions.

**Figure S4 (related to Fig. 5).**
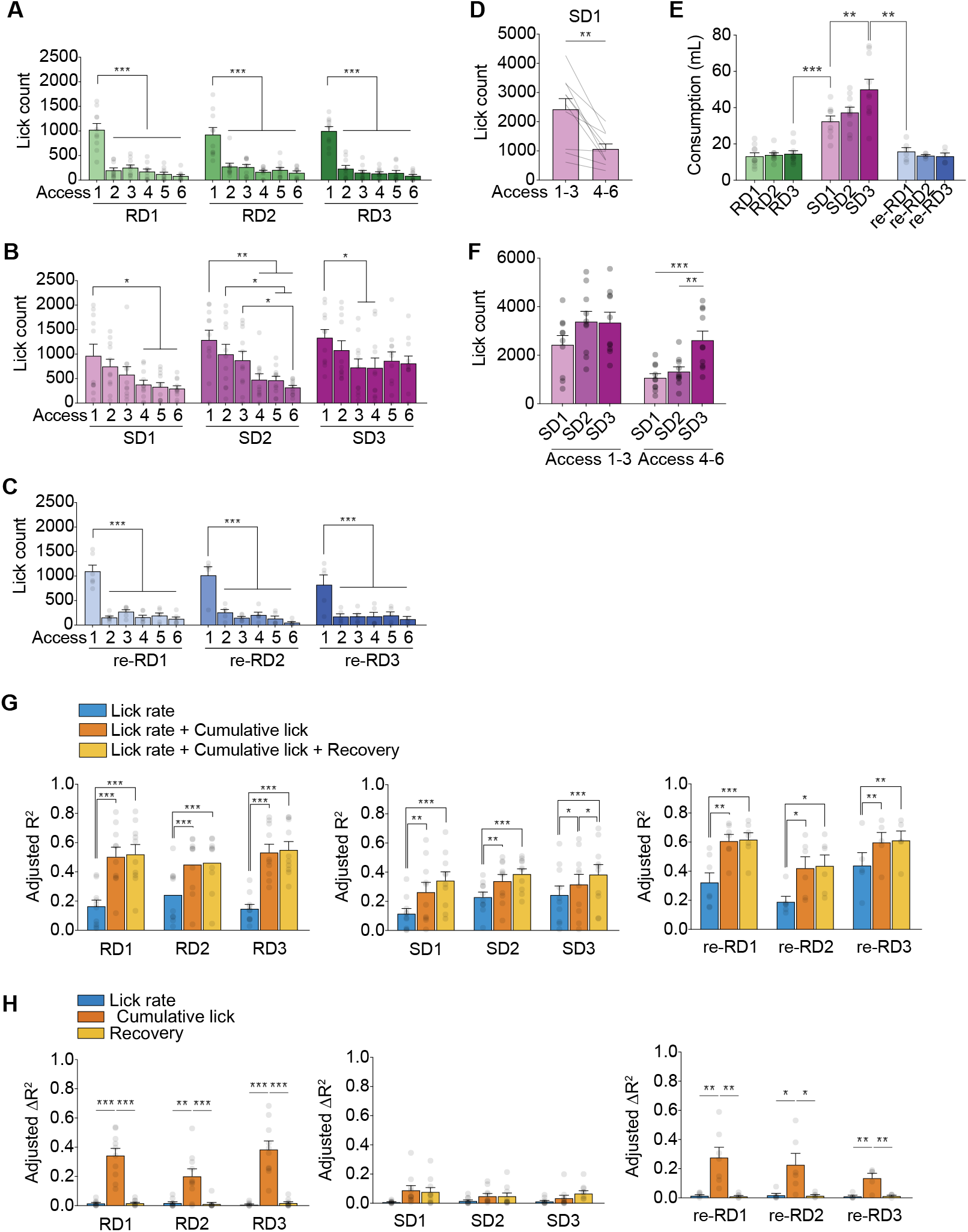
Behavioral adaptation across repeated real and sham drinking sessions and GLM decomposition of MnPO^Agtr1a^ activity. **A-C,** Lick counts during each of the six access periods across repeated real drinking (A; RD1-RD3), sham drinking (B; SD1-SD3), and return to real drinking (C; re-RD1-re-RD3) sessions. **D,** Lick counts during the first versus second half of the first sham-drinking session (SD1), quantified as the summed licks during Accesses 1-3 and Accesses 4-6, respectively. **E,** Total water consumption across repeated RD, SD, and re-RD sessions. **F,** Lick counts during the first three (Accesses 1-3) and last three (Accesses 4-6) access periods across repeated SD sessions. **G,** GLM performance for nested models incorporating lick rate alone, lick rate plus cumulative lick count, or lick rate plus cumulative lick count and post-access recovery, shown across repeated RD, SD, and re-RD sessions. Values indicate adjusted R² for each model. **H,** Unique contributions of the individual lick-rate, cumulative-lick, and post-access-recovery regressors to MnPO^Agtr1a^ activity across repeated RD, SD, and re-RD sessions, quantified as adjusted ΔR². Sample sizes: **A–B, D,** and **F,** n = 10 rats; **C,** re-RD1 n = 6 rats and re-RD2–re-RD3 n = 5 rats; **E,** RD1–RD3 n = 10 rats, SD1 n = 9 rats, SD2–SD3 n = 10 rats, re-RD1 n = 6 rats, and re-RD2–re-RD3 n = 5 rats; **G–H,** RD1–RD3 and SD1–SD3 n = 10 rats, re-RD1 n = 6 rats, and re-RD2–re-RD3 n = 5 rats. \**p* < 0.05, \*\**p* < 0.01, \*\*\**p* < 0.001. All values are reported as mean ± SEM. More detailed statistical information is in **Table S1**.

**Figure S5 (related to Fig. 7).**
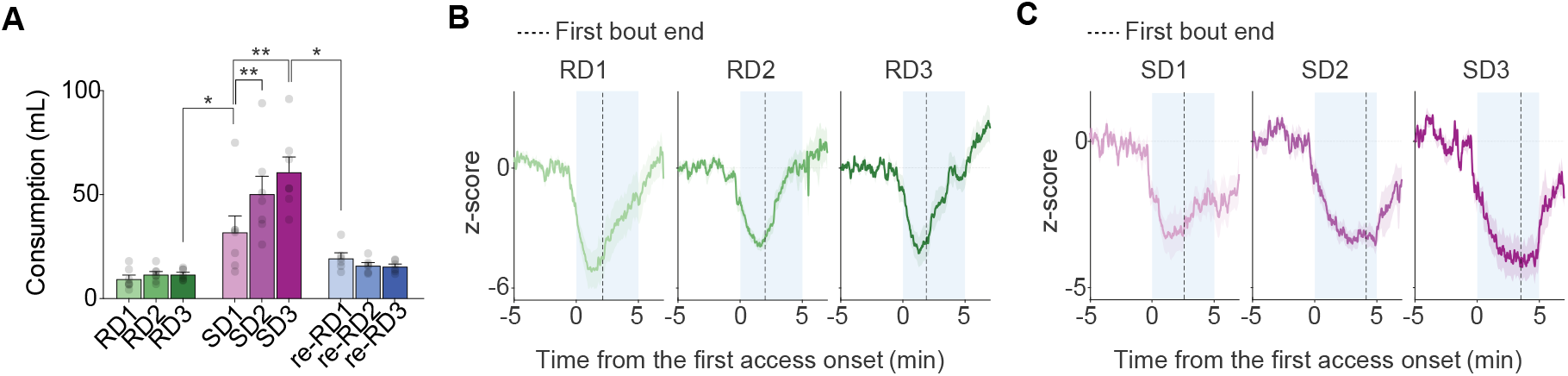
Experience-dependent changes in consumption of hypertonic saline (0.3 M NaCl) and MnPO^Agtr1a^ recovery during hypertonic saline drinking. **A,** Consumption of hypertonic saline (0.3 M NaCl) across repeated real drinking (RD1-RD3), sham drinking (SD1-SD3), and return-to-real-drinking (re-RD1-re-RD3) sessions. **B,** Average MnPO^Agtr1a^ activity traces during repeated real drinking of hypertonic saline (RD1-RD3), aligned to the onset of the first access period. The vertical dashed line indicates the mean time of first lick-bout termination. **C,** Average MnPO^Agtr1a^ activity traces during the first access period across repeated sham-drinking sessions (SD1-SD3), aligned to the onset of the first access period. The vertical dashed line marks the mean time of first lick-bout termination. Sample sizes: **A,** RD1-RD3 and SD1-SD3 n = 8 rats and re-RD1-re-RD3 n = 7 rats; **B-C,** n = 8 rats. \**p* < 0.05, \*\**p* < 0.01. All values are reported as mean ± SEM. More detailed statistical information is in **Table S1**.

## Notes

### Competing Interest Statement

The authors have declared no competing interest.

